# Foxp- and Skor-family proteins control differentiation of Purkinje cells from Ptf1a and Neurogenin1-expressing progenitors in zebrafish

**DOI:** 10.1101/2023.04.06.535843

**Authors:** Tsubasa Itoh, Mari Uehara, Shinnosuke Yura, Jui Chun Wang, Akiko Nakanishi, Takashi Shimizu, Masahiko Hibi

**Affiliations:** Graduate School of Science, Nagoya University, Furo, Chikusa, Nagoya, Aichi 464-8602, Japan

**Keywords:** Ptf1a, Neurogenin1, Skor, Foxp, Purkinje cells, zebrafish

## Abstract

Cerebellar neurons such as Purkinje cells (PCs) and granule cells (GCs) are differentiated from neural progenitors expressing proneural genes. Zebrafish mutants of proneural genes *ptf1a* and *neurogenin1* showed a reduction or loss of PCs, GABAergic interneurons (INs), and reduced expression of GC progenitor genes *atoh1a/b/c*. Lineage tracing revealed that the *ptf1a*-expressing progenitors gave rise to PCs, INs, and a part of GCs in zebrafish. These data indicate that the *ptf1a/neurognin1*-expressing neural progenitors can generate a variety of cerebellar neurons. In this study, we found that genes encoding transcriptional regulators Foxp1b and Foxp4, as well as Skor1b and Skor2, which are reportedly expressed in PCs, were not expressed in *ptf1a*;*neurogenin1* mutants. *foxp1b;foxp4* mutants showed a strong reduction in PCs, while *skor1b;skor2* mutants completely lacked PCs but instead displayed an increase in immature GCs. Misexpression of *skor2* in GC progenitors expressing *atoh1c* suppressed GC fate. These data indicate that Foxp1b/4 and Skor1b/2 function as key transcriptional regulators in the initial step of PC differentiation from *ptf1a/neurogenin1*-expressing neural progenitors, while Skor1b and Skor2 control PC differentiation by suppressing their differentiation into GCs.

## INTRODUCTION

The cerebellum is involved in some forms of skillful movements, motor learning, as well as cognitive and emotional functions. These functions of the cerebellum rely on cerebellar neurons that are conserved among most vertebrates. The cerebellum contains glutamatergic granule cells (GCs) and projection neurons, which are neurons in the deep cerebellar nuclei (DCNs) in mammals or eurydendroid cells (ECs) in teleosts, and GABAergic Purkinje cells (PCs) and interneurons (INs), which include Golgi and stellate cells in both mammals and teleosts, such as zebrafish (Hashimoto and Hibi, 2012; Hibi et al., 2017; Hibi and Shimizu, 2012).

Previous studies in mice revealed that these cerebellar neurons are derived from neural progenitors that express the proneural genes *atoh1* or *ptf1a* (these genes in mice are described as *Atoh1* and *Ptf1a*, but in this study *atoh1* and *ptf1a* will be used for a comparison between animals) (Ben-Arie et al., 1997; Machold and Fishell, 2005; Wang et al., 2005; Wingate, 2005). The *atoh1*-expressing (*atoh1^+^*) neural progenitors are located in the upper rhombic lip (URL, also called the cerebellar rhombic lip) and give rise to projection neurons in DCNs and GCs in the cerebellum (Ben-Arie et al., 1997; Machold and Fishell, 2005; Wang et al., 2005; Wingate, 2005). On the other hand, the *ptf1a*-expressing (*ptf1a^+^*) neural progenitors are located in the ventricular zone (VZ) and give rise to PCs and INs (Hoshino, 2012; Hoshino et al., 2005). In addition to *ptf1a*, proneural genes *Neurogenin1* (*neurog1*) and *Ascl1* are expressed in the VZ of the cerebellum and these proneural gene-expressing neural progenitors were shown to give rise to PCs and INs (Lundell et al., 2009; Sudarov et al., 2011). Expression of *atoh1* (*atoh1a/b/c*) and *ptf1a* genes in the URL and VZ of the cerebellum was also reported for zebrafish (Chaplin et al., 2010; Kani et al., 2010), suggesting similar or identical mechanisms by which proneural genes control the differentiation of cerebella neurons. However, lineage tracing in zebrafish indicated that at least a portion of ECs may be derived from *ptf1a^+^*, suggesting that a slightly different mechanism between mammals and zebrafish may be involved in the differentiation of projection neurons (Kani et al., 2010).

Studies of mouse and zebrafish *atoh1* genes revealed that they are required for the differentiation of GCs (Ben-Arie et al., 1997; Kidwell et al., 2018). Similarly, a mouse *ptf1a* mutant completely lacked PCs and INs (Hoshino et al., 2005) and the zebrafish *ptf1a* mutant showed a reduction – but not loss – of PCs (Itoh et al., 2020), indicating the requirement of *ptf1a* in PC development. The VZ progenitor cells were shown to generate GCs in *ptf1a* mutant mice (Pascual et al., 2007). Ectopic expression of *atoh1* or *ptf1a* in VZ or URL resulted in the generation of glutamatergic and GABAergic neurons, respectively (Yamada et al., 2014), suggesting that expression of *atoh1* and *ptf1a* is sufficient to determine fate of these cell populations. However, it is still not clear whether these proneural genes irreversibly determined the fate of cells in the cerebellum. In the hindbrain region caudal to the cerebellum, the *ptf1a^+^* progenitors give rise to inhibitory neurons in the cochlear nuclei in mice (Fujiyama et al., 2009), excitatory neurons in the inferior olivary nuclei (IO neurons) in both mice and zebrafish (Itoh et al., 2020; Yamada et al., 2007), and crest cells in zebrafish (Itoh et al., 2020), indicating that the *ptf1a^+^* progenitors have the potential to generate neurons other than GABAergic PCs or INs. It was previously shown that the homeodomain transcription factor Gsx2 is involved in fate determination of IO neurons (Itoh et al., 2020). It remains elusive what factors are involved in the differentiation of PCs from the *ptf1a^+^* progenitors in the cerebellum.

Several transcription factors have been shown to be involved in the differentiation of PCs. Forkhead transcription factors Foxp2 and Foxp4 are expressed in PCs in the mouse cerebellum (Ferland et al., 2003; Tam et al., 2011; Tanabe et al., 2012). In the *Foxp2* mutant in mice, even though the specification of PC took place, positioning and dendrite formation of PCs were affected (Shu et al., 2005). siRNA-mediated knockdown of *Foxp4* at a late developmental period resulted in the impairment of PC dendrite formation (Tam et al., 2011). These findings suggest that Foxp-family transcription factors regulate late processes of PC differentiation but are not involved in early differentiation processes. Ski/Sno-family transcriptional co-repressor 2 (Skor2, also known as Corl2) was shown to be expressed in PCs and plays an important role in the differentiation of PCs (Nakatani et al., 2014; Wang et al., 2011). *Skor2* mutant mice exhibited developmental defects in PC development with impaired dendrite arborization, decreased expression of PC marker genes, and increased expression of glutamatergic neuronal genes instead. However, *Skor2* was found to be dispensable for the specification and maintenance of PC fate (Nakatani et al., 2014; Wang et al., 2011). In addition to *Skor2*, *Skor1* is expressed in PCs but its role in PC differentiation remains elusive (Nakatani et al., 2014). Although these transcriptional regulators are involved in some aspects of PC differentiation, it is unclear whether these genes function downstream of Ptf1a and Neurog1. It is also not clear whether they control initial specification of PCs.

Previous RNA-seq analysis of zebrafish cerebellar neurons revealed that *foxp1b/4* and *skor1b/2* are expressed in developing PCs in the zebrafish cerebellum (Takeuchi et al., 2017). In this study, we show that *ptf1a^+^* neural progenitors are capable of generating not only PCs but also INs, ECs, and PCs, and that Foxp1b4 and Skor1b/2 function downstream of Ptf1a and Neurog1 to control differentiation from Ptf1a/Neurog1-expressing neural progenitors into PCs.

## RESULTS

### Ptf1a and Neurog1 are co-expressed in cerebellar VZ progenitors

*ptf1a* is expressed in the cerebellar VZ and involved in the generation of PCs in mice and zebrafish (Hoshino et al., 2005; Kani et al., 2010). PCs are absent in mouse *ptf1a* mutants while PCs are reduced, but not absent, in zebrafish *ptf1a* mutants (Itoh et al., 2020). Lineage tracing in mice suggested that *neurog1* is expressed in the progenitors of PCs in mice (Lundell et al., 2009). Therefore, *neurog1* is a candidate that compensates for the loss of *ptf1a*. We compared the expression of *ptf1a* and *neurog1* by *in situ* hybridization and by using transgenic lines expressing fluorescent proteins (Fig. 1). As reported previously, *ptf1a* transcripts were detected in the cerebellar VZ in early-stage larvae (3 day-post-fertilization [dpf] larvae, Fig. 1A, B), whereas *neurog1* transcripts were barely detected in the cerebellum region (Fig. 1C, D). However, the promoter and enhancer activity of *neurog1* was detected in the cerebellar VZ of *TgBAC(neurog1:EGFP)* (hereafter, named neurog1:EGFP) larvae (Fig. 1G, J). We compared neurog1:GFP-expressing cells with *ptf1a*-expressing (*ptf1a^+^*) cells that were marked by using the Gal4-UAS system with *TgBAC(ptf1a:GAL4-VP16)* and *Tg(UAS:RFP)* (referred to as ptf1a::RFP) (Fig. 1F, I). ptf1a::RFP was detected in the VZ progenitor cells, in the same way that *ptf1a^+^*cells were labeled with *Tg(ptf1a:GFP)* (Fig. S1). We found that some ptf1a::RFP-expressing cells also expressed neurog1:EGFP (Fig. 1E, H), suggesting that at least part of the *ptf1a^+^* neural progenitors also express *neurog1* in the cerebellar VZ.

**Figure 1.**
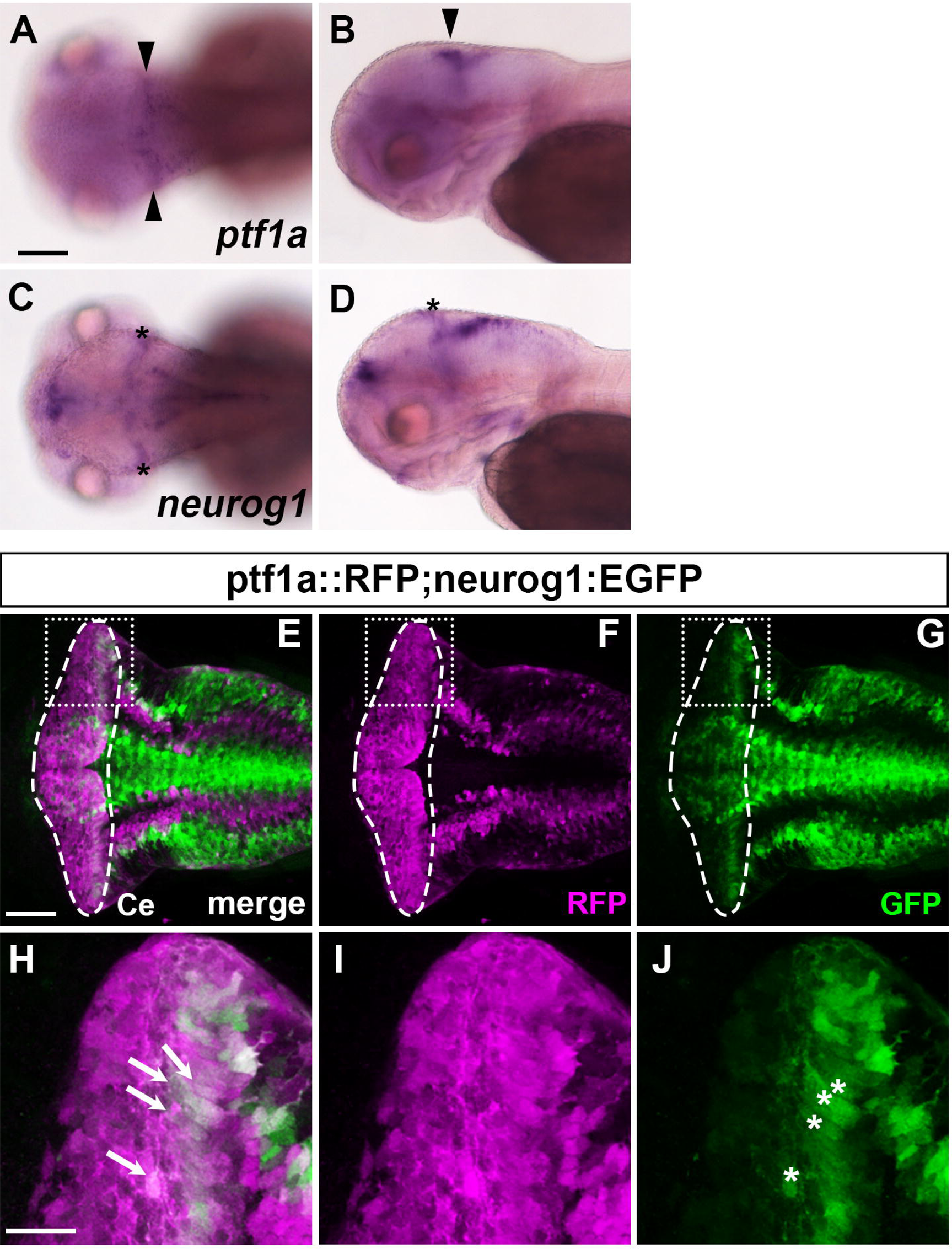
Expression of *ptf1a* and *neurogenin1* in the cerebellum. (A-D) Expression of *ptf1a* (A, B) and *neurogenin1* (*neurog1*, C, D) mRNA at 3-dpf. Transcripts were detected by *in situ* hybridization. Dorsal (A, C) and lateral views (B, D) with anterior to the left. Expression of *ptf1a* in the cerebellar ventricular zone is marked by arrowheads. Expression of *neuurog1*, marked by asterisks, was in the tectum but not the cerebellum. (E-J) Detection of *ptf1a*- and/or *neurog1*-expressing cells using transgenic lines. 5-dpf *Tg(ptf1a:GAL4-VP16); Tg(UAS:RFP); Tg(neurog1:GFP)* larvae (*n*=3) were stained with anti-RFP (magenta) and anti-GFP (green) antibodies. *Tg(ptf1a:GAL4-VP16); Tg(UAS:RFP*) (referred to as ptf1a::RFP). Dorsal views of the rostral hindbrain region, including the cerebellum. The cerebellar region (Ce) is surrounded by a dotted line. (H-J) Higher magnification views of boxes in E, F, G. The ptf1a::RFP and neurog1:GFP double-positive cells are marked by white arrows (H) and the expression of neurog1:GFP*^+^* cells in the cerebellar ventricular zone is indicated by white asterisks (J). Scale bars: 100 μm in A (applies to A-D); 50 μm in E (applies to E-G); 20 μm in H (applies to H-J).

### Ptf1a and Neurog1 cooperate to generate various cerebellar neurons

To reveal the roles of *ptf1a* and *neurog1* in cerebellar neurogenesis, we generated combined mutants of *ptf1a*^Δ*4*^ and *neurog1^hi1059Tg^* (referred to as *neurog1*^-^) alleles, which are supposed to be null alleles (Fig. 2, 3) (Golling et al., 2002; Itoh et al., 2020) and analyzed their phenotypes by marker expression. Whereas *neurog1* mutant larvae had comparable numbers of PCs and INs, which were marked by parvalbmin7 (Pvalb7) and Pax2, compared to wild-type (WT) larvae, *ptf1a* mutants showed a significant reduction in PCs and INs (Fig. 2A-C, E-G, AG, AH, Table 1). The *neurog1* mutation enhanced *ptf1a* mutant phenotypes and *ptf1a;neurog1* double mutant larvae showed an almost complete lack of PCs and INs (Fig. 2D, H, AG, AH). Consistent with this, *ptf1a* mutants showed a reduced expression of genes that are reportedly expressed in zebrafish PCs (Takeuchi et al., 2017), including *foxp1b/4*, *skor1b/2*, *lhx1a*, and *rorb*. *ptf1a:neurog1* mutants displayed lack of expression of these PC genes (Fig. 2I-AF). A similar reduction and loss of crest cells in the anterior hindbrain, which receive GC axons and function in the cerebellum-like structure (Hibi and Shimizu, 2012), was observed in *ptf1a* and *ptf1a:neurog1* mutants, respectively (Fig. S2). These data indicate that *ptf1a* plays a major role in the development of PCs and INs in the cerebellum and crest cells of the rostral hindbrain, but that *neurog1* is not essential for this development, although it has some redundant functions that overlap with those of *ptf1a*.

**Figure 2.**
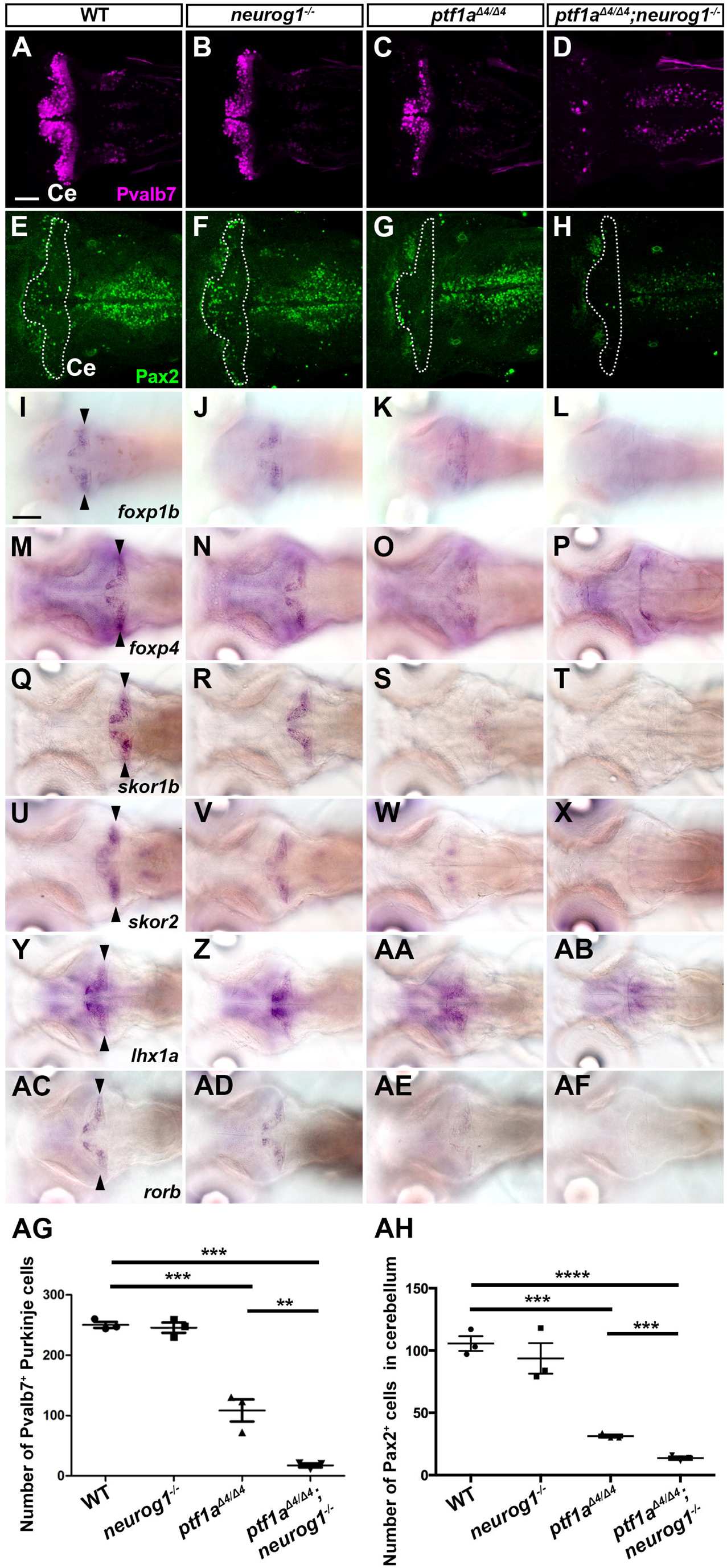
*ptf1a* and *neurog1* are required for the development of GABAergic PCs and INs. (A-Z, AA-AF) Expression of parvalbumin7 (Pvalb7, A-D), Pax2 (E-H), *foxp1b* (I-L), *foxp4* (M-P), *skor1b* (Q-T), *skor2* (U-X), *lhx1a* (Y, Z, AA, AB), and *rorb* (AC-AF) in the cerebellum of 5-dpf wild-type (WT), *neurog1* mutant, *ptf1a* mutant, or *ptf1a;neurog1* double mutant larvae. Immunostaining with anti-Pvalb7 (A-D) and anti-Pax2 antibodies (E-H). *In situ* hybridization (I-Z, AA-AF). Dorsal views with anterior to the left. The cerebellum region (Ce) is surrounded by a dotted line (E-H). Pvalb7, *foxp1b/4*, *skor1b/2*, *lhx1a*, and *rorb* were expressed in PCs (expression of PC genes in the cerebellum is indicated by arrowheads). Pax2 is a marker of GABAergic INs. Note that expression of these PC and IN markers was reduced in *ptf1a* mutant and absent in *ptf1a:neurog1* mutant larvae. The number of examined larvae and larvae showing each expression pattern is described in Table 1. Scale bars: 50 μm in A (applies to A-H); 100 μm in I (applies to I-Z, AA-AF). (AG, AH) Number of Pvalb7^+^ PCs and Pax2^+^ INs in the cerebellum of 5-dpf WT, *neurog1*, *ptf1a*, and *ptf1a;neurog1* mutant larvae was plotted in graphs. \*\**P*<0.01, \*\*\**P*<0.001, \*\*\*\**P*<0.0001 (ANOVA with Tukey’s multiple comparison test). Data are means±SE. with individual values indicated.

**Figure 3.**
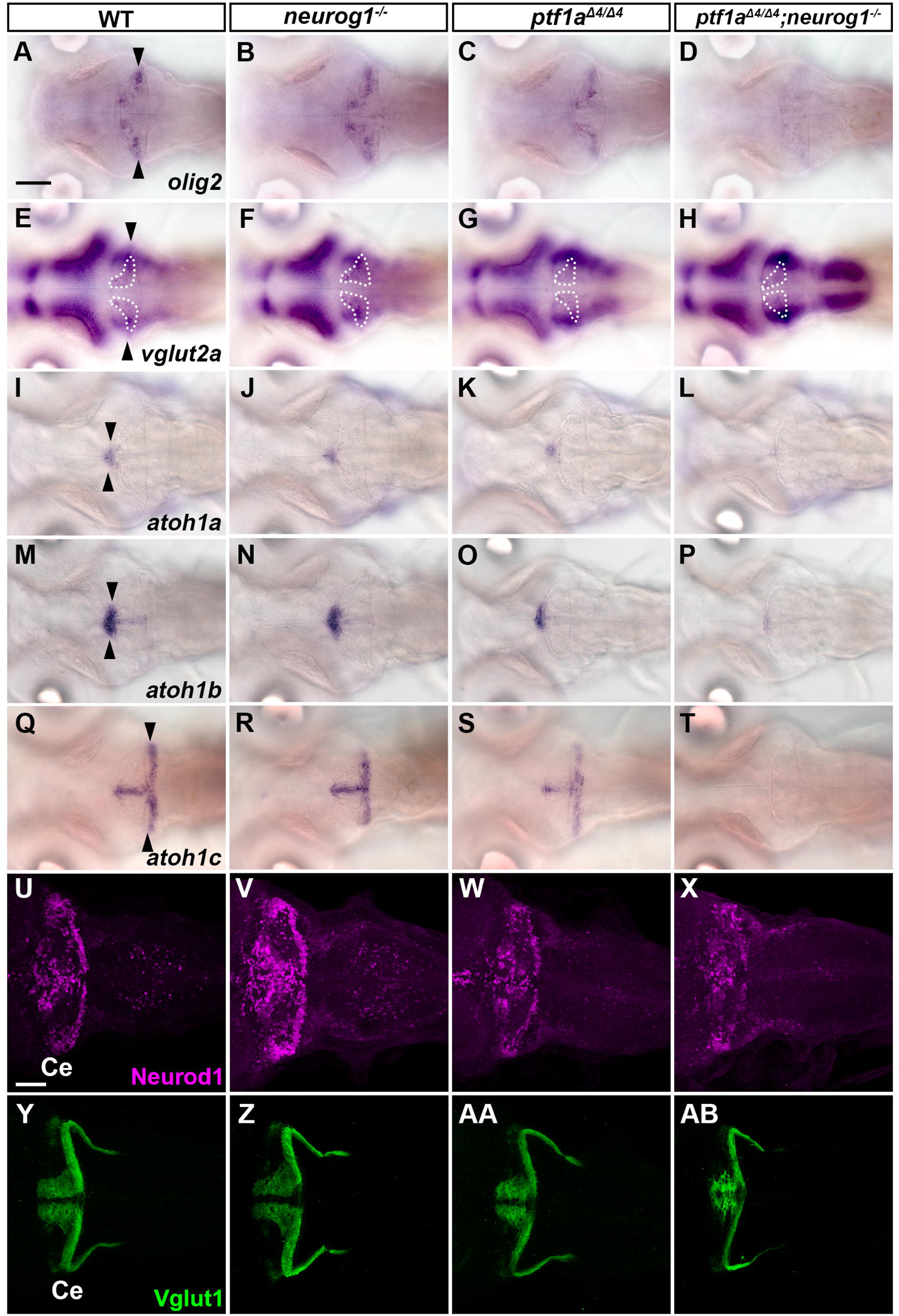
*ptf1a* and *neurog1* are involved in the development of ECs and GCs. (A-T) Expression of *olig2* (A-D), *vglut2a* (E-H), *atoh1a* (I-L), *atoh1b* (M-P), and *atoh1c* (Q-T) in 5-dpf wild-type (WT), *neurog1*, *ptf1a*, and *ptf1a;neurog1* mutant larvae. *olig2* and *vglut2a* were expressed in ECs. *atoh1a/b/c* were expressed in GC progenitors. Expression of these ECs and GCs was reduced in *ptf1a* mutants and absent in *ptf1a;neurog1* mutants. Arrowheads indicate the expression of genes in the cerebellum. Expression of the cerebellum region in E-H is surrounded by a dotted line. (U-Z, AA, AB) Expression of GC markers Neurod1 and Vglut1 in 5-dpf WT, *neurog1*, *ptf1a*, and *ptf1a;neurog1* mutant larvae. Expression pattern of Neurod1 and Vglut1 was affected in *ptf1a* and *ptf1a;neurog1* mutants, but the area of Neurod1-expression domains was variable in *ptf1a* mutants. The number of examined larvae and larvae showing each expression pattern is described in Table 1. Scale bars: 100 μm in A (applies to A-T); 50 μm in U (applies to U-Z, AA, AB).

**Table 1.** Phenotypic summary of *ptf1a* and *neurog1* mutants

| Marker<br>(stage) | Genotype | WT | *neurogl*<sup>-/-</sup> | *ptfla*<sup>Δ4/Δ4</sup> | *ptfla*<sup>Δ4/Δ4</sup><br>;*neurogl*<sup>-/-</sup> |
| --- | --- | --- | --- | --- | --- |
| Proteins |  |  |  |  |  |
| Pvalb7 (5 dpf) |  | +++<br>(n = 3) | +++<br>(n = 3) | ++<br>(n = 3) | +<br>(n = 3) |
| Vglut1 (5 dpf) |  | +++<br>(n = 3) | +++<br>(n = 3) | +++<br>(n = 3) | ++<br>(n = 3) |
| Pax2 (5 dpf) |  | +++<br>(n = 3) | +++<br>(n = 3) | +<br>(n = 3) | -<br>(n = 3) |
| Genes |  |  |  |  |  |
| *atoh1a* (3 dpf) |  | +++<br>(n = 3/3) | +++<br>(n = 3/3) | +++<br>(n = 3/3) | +++<br>(n = 1/1) |
| *atoh1a* (5 dpf) |  | +++<br>(n = 3/3) | +++<br>(n = 3/3) | ++<br>(n = 3/3) | +<br>(n = 3/3) |
| *atoh1b* (3 dpf) |  | +++<br>(n = 3/3) | +++<br>(n = 3/3) | +++<br>(n = 3/3) | +++<br>(n = 3/3) |
| *atoh1b* (5 dpf) |  | +++<br>(n = 3/3) | +++<br>(n = 2/2) | ++<br>(n = 5/5) | +<br>(n = 3/3) |
| *atoh1c* (3 dpf) |  | +++<br>(n = 2/2) | +++<br>(n = 4/4) | +++<br>(n = 4/4) | +++<br>(n = 2/2) |
| *atoh1c* (5 dpf) |  | +++<br>(n = 4/4) | +++<br>(n = 4/4) | +<br>(n = 2/4) | +<br>(n = 1/1) |
| *olig2* (5 dpf) |  | +++<br>(n = 4/4) | +++<br>(n = 4/4) | ++<br>(n = 4/4) | +<br>(n = 4/4) |
| *vglut2a* (5 dpf) |  | +++<br>(n = 4/4) | +++<br>(n = 3/3) | +<br>(n = 4/4) | +<br>(n = 4/4) |
| *foxp1b* (5 dpf) |  | +++<br>(n = 4/4) | +++<br>(n = 4/4) | +<br>(n = 4/4) | -<br>(n = 4/4) |
| *foxp4* (5 dpf) |  | +++<br>(n = 3/3) | +++<br>(n = 3/3) | +<br>(n = 3/3) | -<br>(n = 3/3) |
| *skor1b* (5 dpf) |  | +++<br>(n = 4/4) | +++<br>(n = 4/4) | +<br>(n = 2/2) | -<br>(n = 3/3) |
| *skor2* (5 dpf) |  | +++<br>(n = 3/3) | +++<br>(n = 3/3) | +<br>(n = 2/2) | -<br>(n = 3/3) |
| *lhx1a* (5 dpf) |  | +++<br>(n = 2/2) | +++<br>(n = 3/3) | +<br>(n = 3/3) | -<br>(n = 3/3) |
| *rorb* (5 dpf) |  | +++<br>(n = 3/3) | +++<br>(n = 3/3) | +<br>(n = 3/3) | -<br>(n = 3/3) |

3 or 5-dpf wild-type (WT), *neurogl*, *ptfla*, or *ptfla;neurogl* mutant larvae were fixed and analyzed by immunostaining with anti-Pvalb7 (PC marker), Vglut1 (GC axon marker), or Pax2 (IN marker) antibodies, or by whole mount *in situ* hybridization of riboprobes. Expression levels are indicated by +++, ++, +, and -. +++ indicates expression comparable to that in WT; ++ indicates weak expression, + indicates strongly reduced expression; - indicates little or no expression.

In addition to the PC and IN markers, the expression of *olig2* and *vglut2a* (*slc17a6b*), which were expressed in ECs (Bae et al., 2009; Kani et al., 2010; McFarland et al., 2008), decreased in the *ptf1a* mutant cerebellum, and further decreased in the *ptf1a;neurog1* mutant cerebellum (Fig. 3A-H). Furthermore, the expression of *atoh1a*, *atoh1b*, and *atoh1c*, which were expressed in the GC progenitors (Chaplin et al., 2010; Kani et al., 2010; Kidwell et al., 2018), was not affected at 3 dpf (Fig. S3) but was reduced at 5 dpf in *ptf1a;neurog1* mutant larvae (Fig. 3I-T). *ptf1a;neurog1* mutant larvae had a variable number of cells expressing GC markers Neurod1 and Vglut1 (Slc17a7a) at 5 dpf (Fig. 3X, AB). These data indicate that Ptf1a and Neurog1 are not absolutely essential for the development of glutamatergic ECs and GCs, but are at least partly involved in their development.

### Ptf1a-expressing neural progenitors give rise to a variety of cerebellar neurons

*ptf1a* and *neurog1* are expressed in cerebellar neural progenitors, and their mutation affects the development of various cerebellar neurons. However, this does not necessarily mean that these proneural genes cell-autonomously function in the neural progenitors of cerebellar neurons. To gain some insight, we traced the *ptf1a^+^* cell lineage (Fig. 4, 5). mCherry and CreERT2 were expressed in *ptf1a^+^* cells by using the Gal4-UAS system with *TgBAC(ptf1a:GVP)* and *Tg(UAS-hsp70l:mCherry-T2A-CreERT2)*lines (referred to as ptf1a::mCherry-T2A-CreERT2). A reporter line *TgBAC(gad1b:LOXP-DsRed-LOXP-GFP)* was used to trace GABAergic neurons. In this experiment, when CreERT2 was expressed in *ptf1a^+^* cells and activated with endoxifen, CreERT2 induced recombination of the reporter gene, resulting in the conversion from DsRed to GFP expression in GABAergic neurons. GFP-expressing (GFP^+^) cells are GABAergic neurons derived from *ptf1a^+^* neural progenitors. In the absence of CreERT2 expression, only a small number of GFP^+^ cells was observed (Fig. 4A-F, S), whereas a significant number of GFP^+^ cells was observed in the cerebellum in the presence of ptf1a::mCherry-T2A-CreERT2 (Fig. 4I, L). Endoxifen treatment increased the number of GFP^+^ cells (Fig. 4O, R, S). The likely reason for the expression of GFP in the absence of endoxifen treatment is due to the strong expression of CreERT2 and leakiness of the reporter. The increase in GFP^+^ cells by endoxifen at 2 dpf, when the expression domains of *ptf1a* and *atoh1a* are completely separated from each other in the cerebellum (Kani et al., 2010), indicates that most if not all GFP^+^ cells are derived from neural progenitors expressing *ptf1a* but not *atoh1* at 2 dpf. There were two types of GFP^+^ cells: Pvalb7-expressing (Pvalb7^+^) and Pvalb7-negative (Pvalb7^-^) cells (Fig. 4P-R), which correspond to PCs and INs, respectively. Both GFP^+^ Pvalb7^+^ and GFP^+^ Pvalb7^-^ cells in 5 dpf larvae harboring CreERT2 and the reporter were increased by endoxifen treatment (Fig. 4T, U), indicating that the increased PCs and INs were derived from *ptf1a^+^*neural progenitors at 2 dpf.

**Figure 4.**
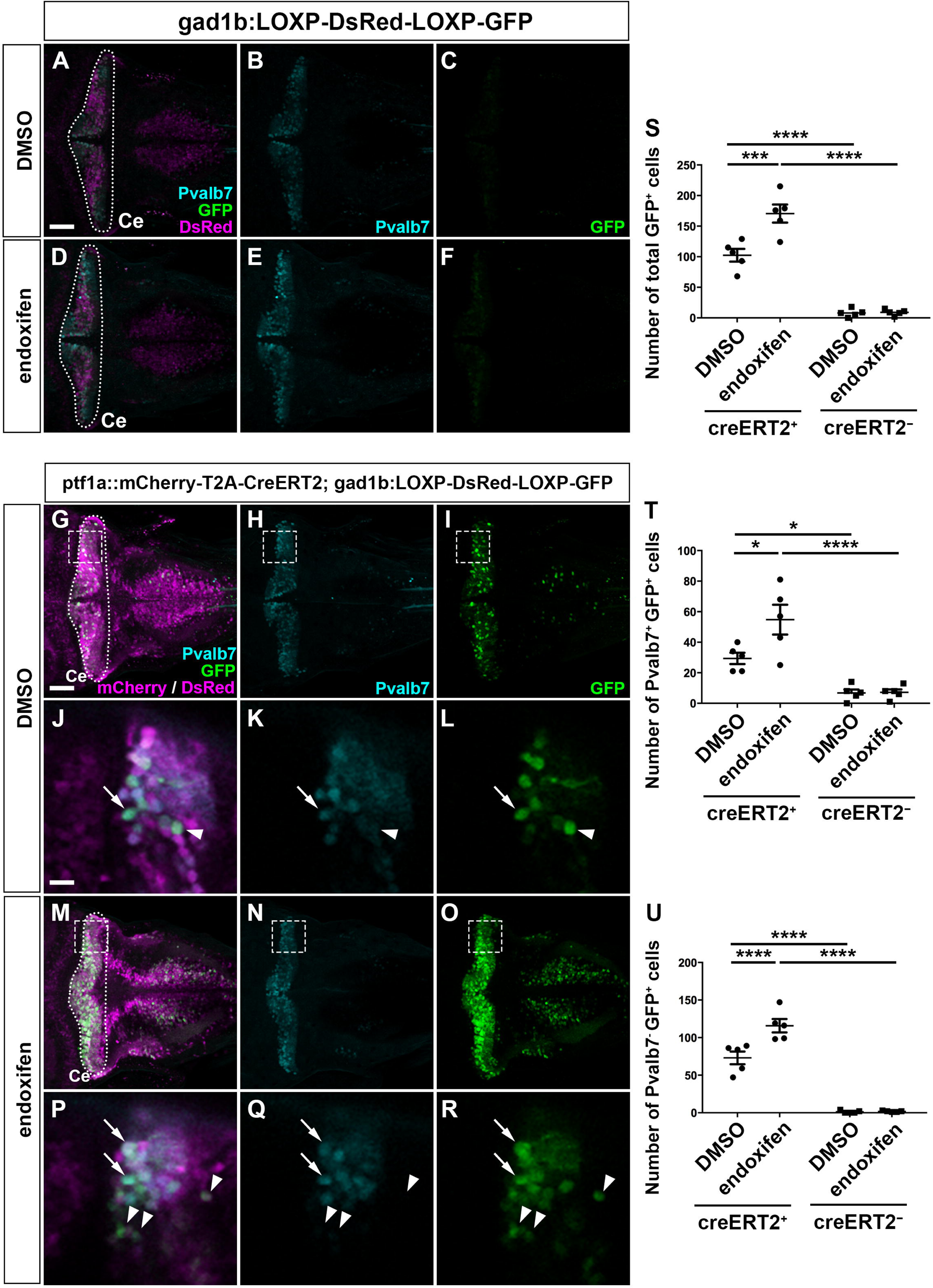
GABAergic PCs and INs were derived from Ptf1a-expressing neural progenitors. (A-F) Expression of Pvalb7 and GFP in 5-dpf *TgBAC(gad1b:LOXP-DsRed-LOXP-GFP)* larvae that were treated with DMSO (control, *n*=5, A-C) or endoxifen (*n*=5, D-F) at 2-dpf. (G-R) Expression of Pvalb7 and GFP in 5-dpf *TgBAC(ptf1a:Gal4-VP16); Tg(UAS-hsp70l:mCherry-T2A-CreERT2); TgBAC(gad1b:LOXP-DsRed-LOXP-GFP)* larvae that were treated with DMSO (*n*=5, G-L) or endoxifen (*n*=5, M-R) at 2-dpf. The larvae were stained with anti-Pvalb7 (cyan), anti-RFP (magenta), and anti-GFP (green) antibodies. Dorsal views with anterior to the left. The cerebellum region (Ce) is surrounded by a dotted line. (J-L, P-R) Higher magnification views of boxes in (G-I, M-O). Arrows and arrowheads indicate Pvalb7^+^ GFP^+^ cells (PCs) and Pvalb7^-^ GFP^+^ cells (INs). Scale bars: 50 μm in A (applies to A-F); 50 μm in G (applies to G-I, M-O); 10 μm in J (applies to J-L, P-R). (S-U) Total number of GFP^+^ cells (S), Pvalb7^+^ GFP^+^ cells (T), and Pvalb7^-^ GFP^+^ cells (U). \**P*<0.05, \*\*\**P*<0.001, \*\*\*\**P*<0.0001 (two-way ANOVA followed by Bonferroni multiple comparisons). Data are means±SE with individual values indicated.

**Figure 5.**
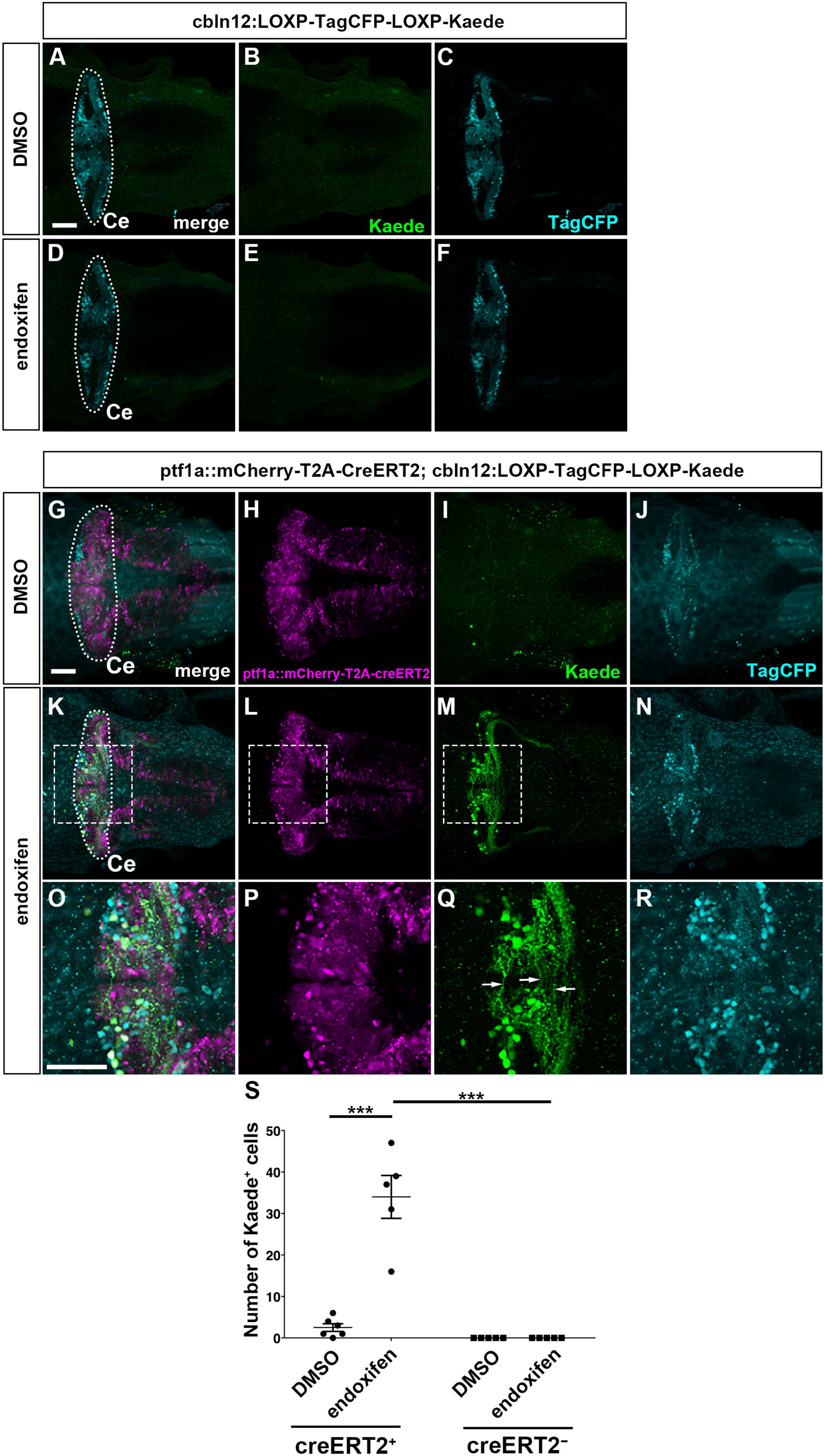
Some GCs were also derived from Ptf1a-expressing neural progenitors. (A-F) Expression of TagCFP (cyan) and Kaede (green) in 5-dpf *Tg(cbln12:LOXP-TagCFP-LOXP-Kaede)* larvae that were treated with DMSO (control, *n*=5, A-C) or endoxifen (*n*=5, D-F) at 2-dpf. (G-R) Expression of TagCFP and Kaede in 5-dpf *TgBAC(atoh1c:Gal4FF); Tg(UAS-hsp70l:RFP-T2A-CreERT2); Tg(cbln12:LOXP-TagCFP-LOXP-Kaede)* larvae that were treated with DMSO (*n*=6, G-J) or endoxifen (*n*=5, K-R) at 2 dpf. The larvae were stained with anti-TagCFP (cyan), anti-RFP (magenta), and anti-Kaede (green) antibodies. Dorsal views with anterior to the left. The cerebellum region (Ce) is surrounded by a dotted line. (O-Q) Higher magnification views of boxes in (K-M). Arrows indicate parallel fibers of GCs. Scale bars: 50 μm in A (applies to A-F); 50 μm in G (applies to G-N); 50 μm in O (applies to O-R). (S) Number of Kaede^+^ cells. \*\*\**P*<0.001 (two-way ANOVA followed by Bonferroni multiple comparisons). Data are means±SE with individual values indicated.

We further examined the GC lineage derived from *ptf1a^+^* neural progenitors by using a reporter line *Tg(cbln12:LOXP-TagCFP-LOXP-Kaede)* (Fig. 5C), which expresses TagCFP in GCs in a *cbln12* promoter-dependent manner (Dohaku et al., 2019). In this experiment, the expression and activation of CreERT2 induced recombination of the reporter gene, resulting in a conversion from TagCFP to Kaede expression in GCs. Kaede-expressing (Kaede^+^) cells are GCs derived from *ptf1a^+^* neural progenitors. Kaede was barely detected in larvae with only the reporter gene (Fig. 5B, E), and in larvae with both CreERT2 and reporter genes but no endoxifen treatment (Fig. 5I), indicating that this reporter had very low leakiness. Endoxifen treatment at 2 dpf resulted in the appearance of Kaede^+^ cells that extended typical parallel fibers (Fig. 5M, Q). These data indicate that a portion of GCs in the cerebellum was derived from *ptf1a^+^* neural progenitors in zebrafish. Considering the data for both *ptf1a* and *neurog1* mutants, Ptf1a/Neurog1-expressing neural progenitors are capable of generating a variety of cerebellar neurons.

### Foxp1b/4 and Skor1b/2 function downstream of Ptf1a and Neurog1 in differentiating PCs

There should be regulators that control the specification and/or differentiation of PCs from Ptf1a/Neurog1-expressing neural progenitors. We previously identified genes that were preferentially expressed in larval PCs (Takeuchi et al., 2017). Among them, we focused on genes encoding transcriptional regulators. *foxp*-family *foxp1b*, *foxp4* and *skor*-family *skor1b* and *skor2* were expressed in PCs in 5 dpf WT larvae but were absent in *ptf1a;neurog1* double mutant larvae (Fig. 2L, P, T, X), suggesting that these genes function downstream of Ptf1a and Neurog1. We generated antibodies against Foxp1b, Skor1b, and Skor2 and used them to analyze their expression by co-immunostaining with anti-Pvalb7 antibody. Foxp1b was detected in Pvalb7^+^ PCs as well as in Pvalb7^-^ cells in the cerebellum of WT larvae (Fig. 6A-F), but was not observed in the cerebellum of *foxp1b* mutant larvae (Fig. 6G-L) (the *foxp1b* mutant is described below). Foxp1b was also detected in the nucleus of PCs in the WT adult cerebellum, but not in the *foxp1b* mutant cerebellum (Fig. 6O, R). Both Skor1b and Skor2 were detected in Pvalb7^+^ PCs and Pvalb7^-^ cells in the larval but not adult cerebellum (Fig. 6S-X, AE-AJ), but were not observed in *skor1b* and *skor2* mutant larvae (Fig. 6Y-AD, AK-AP) (*skor1b* and *skor2* mutants are described below). Although the possibility that Foxp1b, Skor1b and Skor2 are expressed in non-PC lineage cells of the cerebellum cannot be completely excluded, the data suggest that these proteins are expressed in PC lineage cells before PCs become fully differentiated.

**Figure 6.**
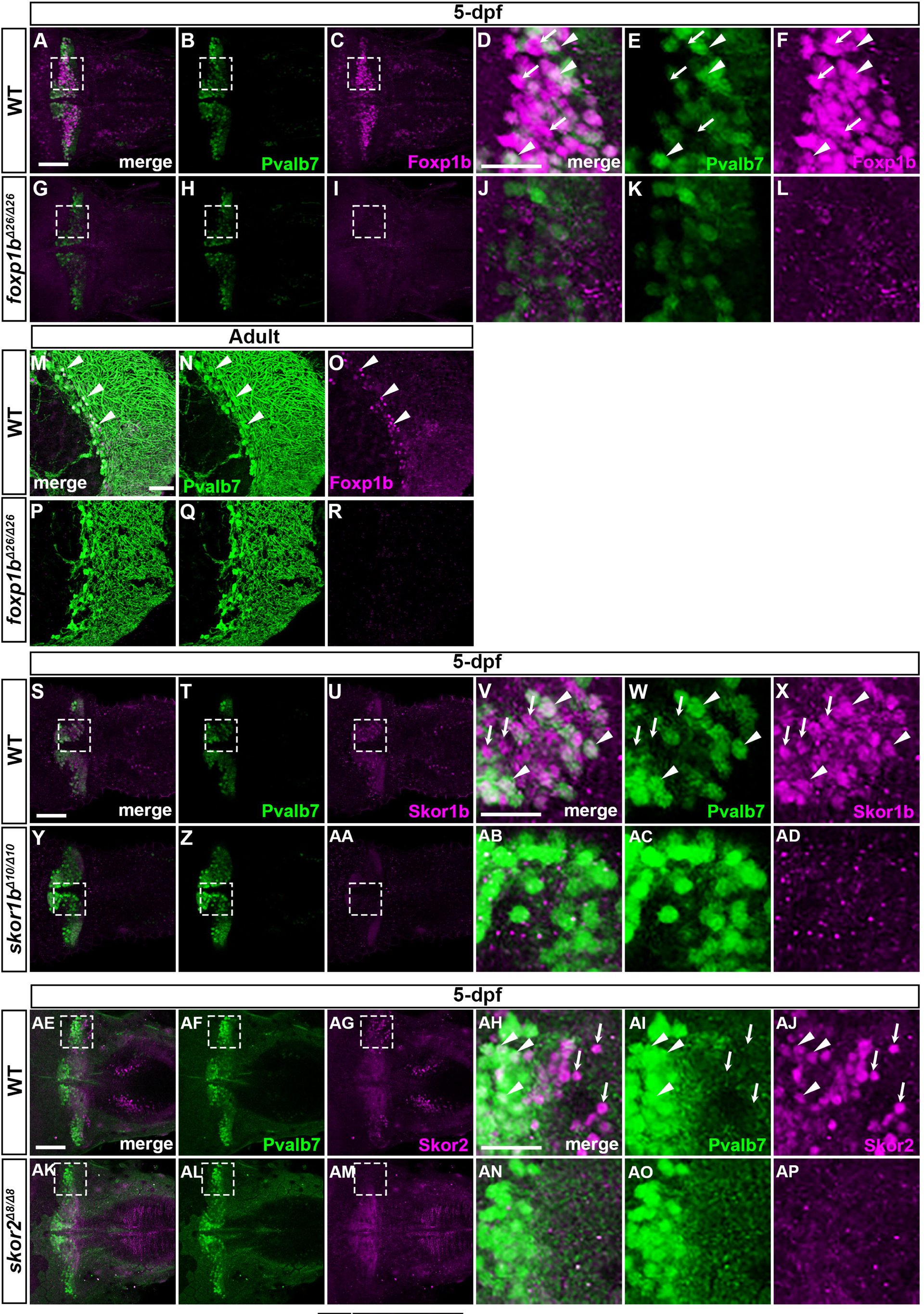
Foxp1b, Skor1b, and Skor2 were expressed in differentiating and differentiated PCs. (A-R) Localization of Foxp1b. 5-dpf WT (*n*=3, A-F) and *foxp1b* mutant larvae (*n*=3, G-L), and adult wild-type (WT) (*n*=2, M-O) and *foxp1b* mutant (*n*=2, P-R) cerebellum sections were immuno-stained with anti-Foxp1b (magenta) and ant-Pvalb7 antibodies (green). Dorsal views with anterior to the left (A-L) and sagittal sections (M-R). (D-F, J-L) Higher magnification views of boxes in (A-C, G-I). Arrowheads and arrows indicate examples of Foxp1b^+^ Pvalb7^+^ cells and Foxp1b^+^ Pvalb7^-^ cells, respectively (D-F, M-O). (S-AQ) Localization of Skor1b and Skor2. (S-AD) 5-dpf WT (*n*=3, S-X) and *skor1b* mutant larvae (*n*=3, Y-AD) were immunostained with anti-Skor1b (magenta) and anti-Pvalb7 antibodies (green). (AE-AP) 5-dpf WT (*n*=2, AE-AJ) and *skor2* mutant larvae (*n*=2, AK-AP) were immunostained with anti-Skor2 (magenta) and anti-Pvalb7 antibodies (green). Dorsal views with anterior to the left. (V-X, AB-AD, AH-AJ, AN-AP) Higher magnification views of boxes in (S-U, Y-AA, AE-AG, AK-AM). Scale bars: 50 μm in A (applies to A-C, G-I); 50 μm in D (applies to D-F, J-L); 50 μm in M (applies to M-R); 50 μm in S (applies to S-U, Y-AA); 50 μm in V (applies to V-X, AB-AD); 50 μm in AE (applies to AE-AG, AK-AM); 50 μm in AH (applies to AH-AJ, AN-AP). Arrowheads indicate examples of Skor1b^+^ Pvalb7^+^ cells (V-X) and Skor2^+^ Pvalb7^+^ cells (AH-AJ). Arrows indicate examples of Skor1b^+^ Pvalb7^-^ cells (V-X) and Skor2^+^ Pvalb7^-^ cells (AH-AJ).

### Foxp1b/4 and Skor1b/2 are required for the differentiation of PCs

We generated mutants of *foxp1b/4* and *skor1b/2* using the CRISPR/Cas9 method (Fig. S4). The *foxp1b* and *foxp4* mutants harbor 26- and 7-bp deletions in exon 14 of *foxp1b* and exon 7 of *foxp4*, respectively, that introduce a premature stop codon. The putative mutant Foxp1b and Foxp4 proteins lacked the DNA-binding forkhead domain, so these mutations are likely null alleles. The *skor1b* and *skor2* mutants harbor 10- and 8-bp deletions in exon 1 of *skor1b* and exon 2 of *skor2*, respectively, that introduce a premature stop codon. Although the functional domains of Skor proteins were not well understood, the putative mutant Skor1b and Skor2 proteins lacked the protein from the c-Ski SMAD binding domain to the carboxy-terminus, so these mutations are potentially null alleles.

Single mutant larvae of *foxp1b* or *foxp4* showed a slight reduction in the expression of Pvalb7, zebrin II (encoded by *aldolase Ca* gene), carboxy anhydrase 8 (Ca8), or *rorb* in the cerebellum (Fig. 7A-C, E-G, I-K, U-W). The *foxp1b;foxp4* double mutant displayed a more severe reduction in these PC markers (Fig. 7D, H, L, X). After counting the number of Pvalb7^+^ PCs in the mutants, it was confirmed that PCs were slightly reduced in *foxp1b* and *foxp4* single mutants compared to WT, but were more severely reduced in *foxp1b;foxp4* double mutants (Fig. 7AO). In contrast, expression of the GC markers Neurod1 and Vglut1, the EC markers *olig2* and *vglut2a*, and the IN marker *pax2a* was not affected in either single or double mutants (Fig. 7M-T, Table 2). These data suggest that Foxp1b and Foxp4 function partially redundantly in PC differentiation; Foxp1b and Foxp4 are required for the proper differentiation of PCs but not GCs, ECs, or INs, in the cerebellum.

**Figure 7.**
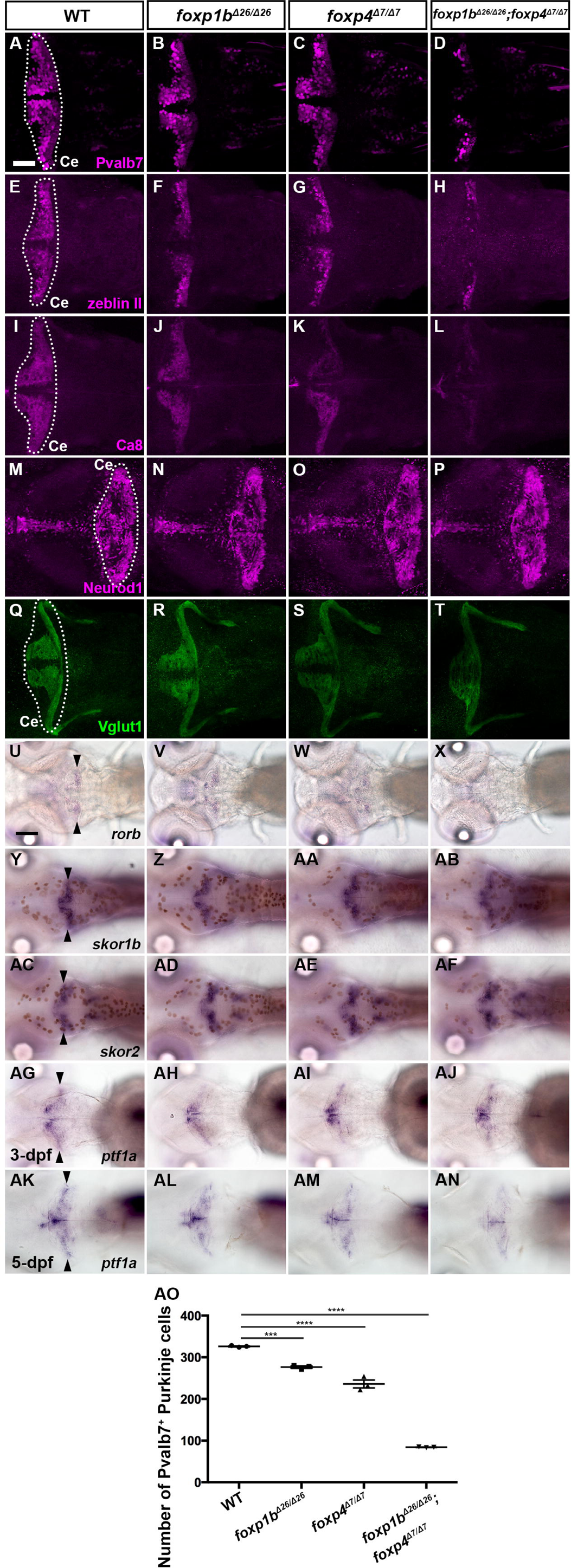
Phenotypes of *foxp1b* and *foxp4* mutants. (A-T) Expression of PC markers Pvalb7, Zebrin II, and Ca8, and GC markers Neurod1, and Vglut1 in 5-dpf wild-type (WT), *foxp1b*, *foxp4*, and *foxp1b;foxp4* mutant larvae. (U-AB, AG-AJ) Expression of *rorb*, *skor1b*, *skor2* and *ptf1a* in 5-dpf WT, *foxp1b*, *foxp4*, and *foxp1b;foxp4* mutant larvae. (AG-AJ) Expression of *ptf1a* in 3-dpf WT, *foxp1b*, *foxp4*, and *foxp1b;foxp4* mutant larvae. Data of immunostaining (A-L) and *in situ* hybridization (U-AN). Dorsal views with anterior to the left. The cerebellum region (Ce) is surrounded by a dotted line. Arrowheads indicate expression of genes in the cerebellum. The number of examined larvae and larvae showing each expression pattern is shown in Table 2. Scale bars: 50 μm in A (applies to A-T); 100 μm in U (applies to U-Z, AA-AN). (AO) Number of Pvalb7^+^ PCs in the cerebellum of 5-dpf WT, *foxp1b*, *foxp4*, and *foxp1b;foxp4* mutant larvae. \*\*\**P*<0.001, \*\*\*\**P*<0.0001 (ANOVA with Tukey’s multiple comparison test). Data are means±SE with individual values indicated.

**Table 2.**
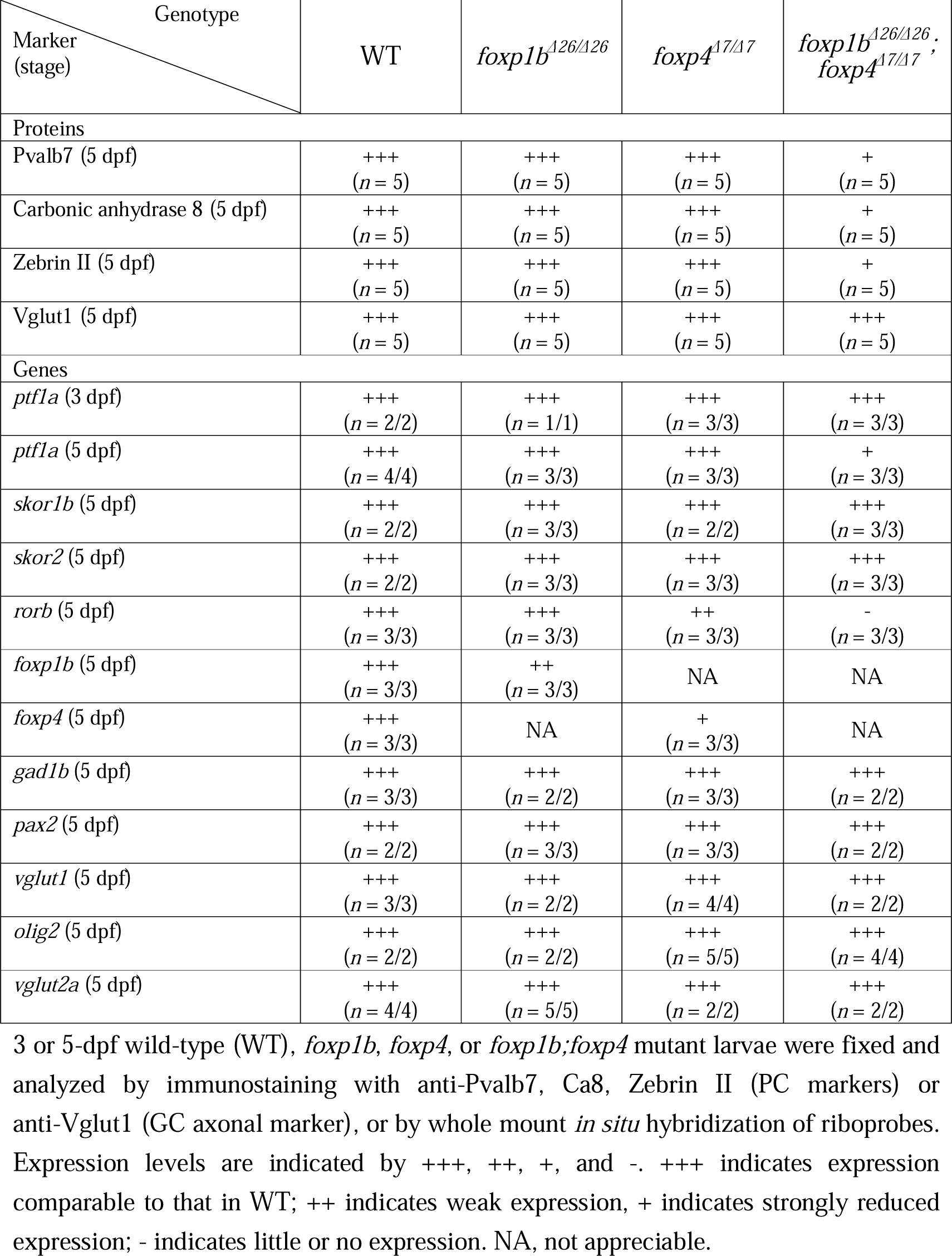
Phenotypes of *foxp1b* and *foxp4* mutants

Single mutant *skor1b* and *skor2* larvae did not show reduced expression of the PC markers compared to WT larvae (Fig. 8A-C, E-G, I-K, Q-S, AK), whereas *skor1b;skor2* double mutant larvae showed a complete loss of expression of the PC markers (Pvalb7, Zebin II, Ca8, *rorb*, Fig. 8D, H, L, T, AK). *skor1b;skor2* larvae showed expression of the GC markers Neurod1 and Vglut1 although their expression pattern was affected (Fig. 8P, 9, described below). The expression of EC markers *olig2* and *vglut2a* and IN marker *pax2a* was not affected in either *skor1b*, *skor2* single or *skor1b;skor2* double mutants (Table 3). These data indicate that Skor1b and Skor2 function redundantly and are essential for the differentiation of PCs in the cerebellum.

**Figure 8.**
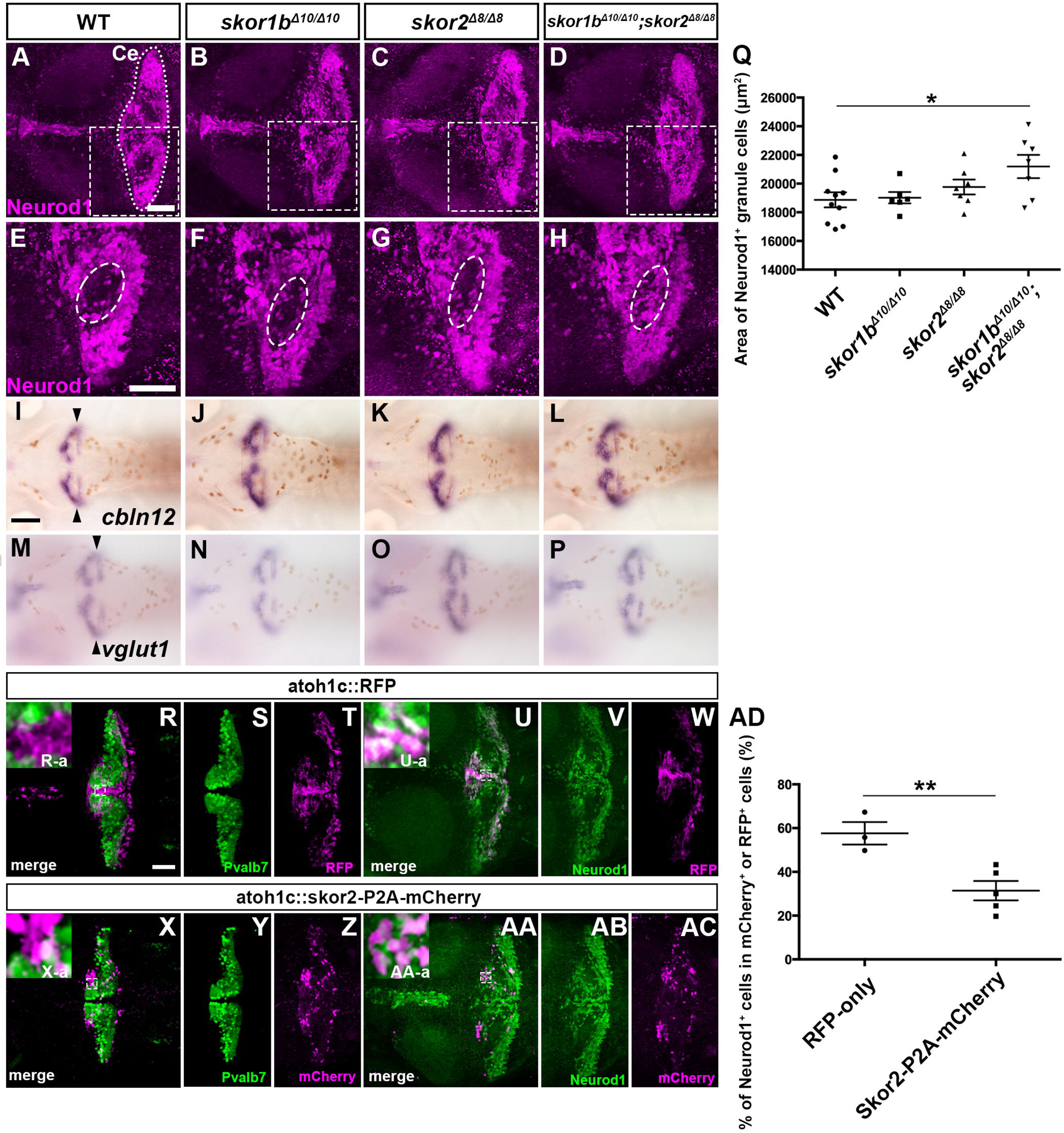
Phenotypes of *skor1b* and *skor2* mutants. (A-P) Expression of PC markers Pvalb7, Zebrin II, and Ca8, and a GC marker Vglut1 in 5-dpf wild-type (WT), *skor1b*, *skor2*, and *skor1b;skor2* mutant larvae. (Q-AB, AG-AJ) Expression of *rorb*, *foxp1b*, *foxp4* and *ptf1a* in 5-dpf WT, *skor1b*, *skor2*, and *skor1b;skor2* mutant larvae. (AC-AF) Expression of *ptf1a* in 3-dpf WT, *skor1b*, *skor2*, and *skor1b;skor2* mutant larvae. Data of immunostaining (A-P) and *in situ* hybridization (Q-AJ). Dorsal views with anterior to the left. Dorsal views with anterior to the left. The cerebellum region (Ce) is surrounded by a dotted line. Arrowheads indicate expression of genes in the cerebellum. Arrows indicate expression of *foxp1b* and *foxp4* in GCs in caudal and rostral parts of the cerebellum (X, AB). The number of examined larvae and larvae showing each expression pattern is shown in Table 3. Scale bars: 50 μm in A (applies to A-P); 100 μm in Q (applies to Q-AJ). (AK) Number of Pvalb7^+^ PCs in the cerebellum of 5-dpf WT, *skor1b*, *skor2*, and *skor1b;skor2* mutant larvae. \*\*\*\**P*<0.0001 (ANOVA with Tukey’s multiple comparison test). Data are means±SE with individual values indicated.

**Figure 9.**
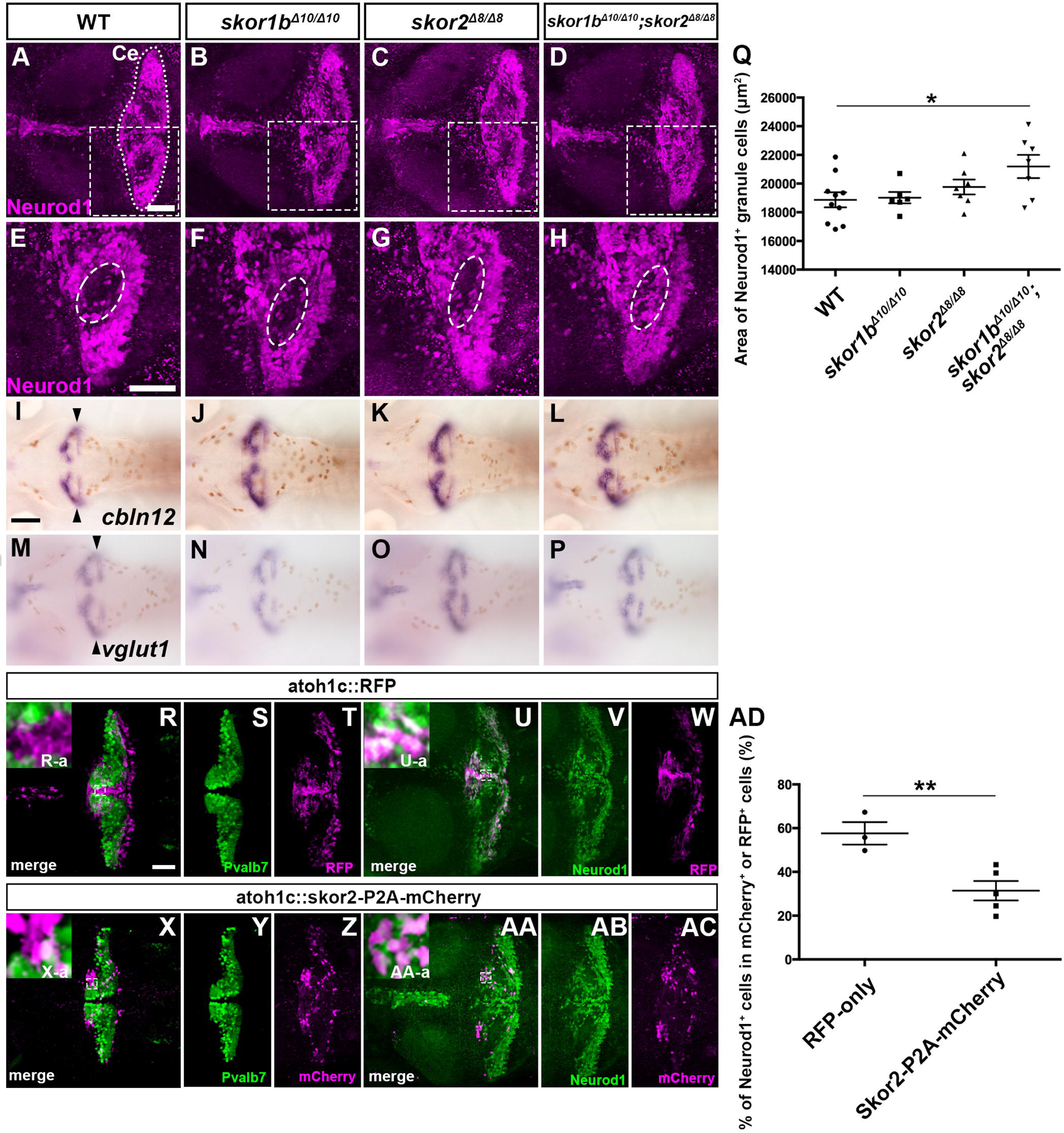
Suppression of granule cell fates by Skor1b/2 and Foxp1b/4. (A-H) Expression of Neurod1 in 5-dpf wild-type (WT), *skor1b*, *skor2*, and *skor1b;skor2* mutant larvae. The cerebellum region is surrounded by a dotted line. (E-H) Higher magnification views of boxes in A-D. Neurod1-expressing GCs were absent in the central areas of the cerebellum (marked by dotted circles) of WT, *skor1b* and *skor2* mutant larvae, but present in the entire cerebellum of *skor1b;skor2* mutant larvae. (I-P) Expression of mature GC marker genes *cbln12* and *vglut1* in the cerebellum. The expression of *cbln12* and *vglut1* was not affected in *skor1b*, *skor2*, and *skor1b;skor2* mutant larvae. (Q) Area of Neurod1^+^ GCs in the cerebellum of 5-dpf WT, *skor1b*, *skor2*, and *skor1b;skor2* mutants. \**P*<0.05 (ANOVA with Tukey’s multiple comparison test). (R-AC) Misexpression of Skor2 in *atoh1c*-expressing neural progenitors. 5-dpf *Tg(atoh1c:Gal4FF);Tg(UAS-hsp70l:RFP)* or *Tg(atoh1c:Gal4FF);Tg(UAS:HA-skor2-P2A-mCherry)* larvae, which express RFP or Skor2/mCherry in the GC lineage, were immunostained with anti-RFP/mCherry (magenta), and Pvalb7 (green, R-T, X-Z) or Neurod1 (green, U-W, AA-AC) antibodies. Dorsal views with anterior to the left (A-P, R-AC). (R-a, U-a, X-a, AA-a) Higher magnification views of boxed in R, U, X, and AA. Scale bars: 50 μm in A (applies to A-D); 50 μm in E (applies to E-H); 100 μm in I (applies to I-P); 100 μm in R (applies to R-AC). (AD) Ratios of Neurod1^+^ cells in mCherry^+^ cells are indicated. \*\**P*<0.01 (Student t-test). Data are means±s.e.m. with individual values indicated (Q, AD).

**Table 3.** Phenotypes of *skor1b* and *skor2* mutants

| Marker<br>(stage) \ Genotype | WT | *skor1b*<sup>Δ10/Δ10</sup> | *skor2*<sup>Δ8/Δ8</sup> | *skor1b*<sup>Δ10/Δ10</sup>;<br>*skor2*<sup>Δ8/Δ8</sup> |
| --- | --- | --- | --- | --- |
| **Proteins** |  |  |  |  |
| Pvalb7 (5 dpf) | +++<br>(n = 5) | +++<br>(n = 5) | +++<br>(n = 5) | -<br>(n = 5) |
| Ca 8 (5 dpf) | +++<br>(n = 5) | +++<br>(n = 5) | +++<br>(n = 5) | -<br>(n = 5) |
| Zebrin II (5 dpf) | +++<br>(n = 5) | +++<br>(n = 5) | +++<br>(n = 5) | -<br>(n = 5) |
| Vglut1 (5 dpf) | +++<br>(n = 3) | +++<br>(n = 3) | +++<br>(n = 3) | +++<br>(n = 3) |
| Neurod1 (5 dpf) | +++<br>(n = 10) | +++<br>(n = 6) | +++<br>(n = 7) | ++++<br>(n = 7) |
| **Genes** |  |  |  |  |
| *ptfla* (3 dpf) | +++<br>(n = 3/3) | +++<br>(n = 3/3) | +++<br>(n = 3/3) | +++<br>(n = 2/2) |
| *ptfla* (5 dpf) | +++<br>(n = 1/1) | +++<br>(n = 2/2) | +++<br>(n = 2/2) | +++<br>(n = 2/2) |
| *rorb* (5 dpf) | +++<br>(n = 3/3) | ++<br>(n = 2/3) | +<br>(n = 3/3) | -<br>(n = 3/3) |
| *skor1b* (5 dpf) | +++<br>(n = 3/3) | ++<br>(n = 2/3) | NA | NA |
| *skor2* (5 dpf) | +++<br>(n = 3/3) | NA | ++++<br>(n = 3/3) | NA |
| *foxp1b* (5 dpf) | +++<br>(n = 2/2) | +++<br>(n = 2/2) | +++<br>(n = 2/2) | ++ *1<br>(n = 2/2) |
| *foxp4* (5 dpf) | +++<br>(n = 3/3) | +++<br>(n = 3/3) | +++<br>(n = 3/3) | +++ *2<br>(n = 2/2) |
| *atoh1a* (3 dpf) | +++<br>(n = 3/3) | +++<br>(n = 2/2) | +++<br>(n = 3/3) | +++<br>(n = 3/3) |
| *atoh1a* (5 dpf) | +++<br>(n = 1/1) | +++<br>(n = 1/1) | +++<br>(n = 1/1) | +++<br>(n = 1/2) |
| *atoh1b* (3 dpf) | +++<br>(n = 3/3) | +++<br>(n = 3/3) | +++<br>(n = 3/3) | +++<br>(n = 1/1) |
| *atoh1b* (5 dpf) | +++<br>(n = 1/1) | +++<br>(n = 1/1) | +++<br>(n = 1/1) | +++<br>(n = 1/1) |
| *atoh1c* (3 dpf) | +++<br>(n = 3/3) | +++<br>(n = 3/3) | +++<br>(n = 3/3) | +++<br>(n = 2/2) |
| *atoh1c* (5 dpf) | +++<br>(n = 2/2) | +++<br>(n = 1/1) | +++<br>(n = 1/1) | +++<br>(n = 3/3) |
| *gad1b* (5 dpf) | +++<br>(n = 2/2) | +++<br>(n = 3/3) | +++<br>(n = 3/3) | +++<br>(n = 1/1) |
| *pax2* (5 dpf) | +++<br>(n = 3/3) | +++<br>(n = 1/1) | +++<br>(n = 3/3) | +++<br>(n = 3/3) |
| *vglut1* (5 dpf) | +++<br>(n = 3/3) | +++<br>(n = 3/3) | +++<br>(n = 3/3) | +++<br>(n = 3/3) |
| *cbln12* (5 dpf) | +++<br>(n = 1/1) | +++<br>(n = 2/2) | +++<br>(n = 3/3) | +++<br>(n = 4/4) |
| *pax6a* (5 dpf) | +++<br>(n = 3/3) | +++<br>(n = 3/3) | +++<br>(n = 3/3) | +++<br>(n = 1/1) |
| *reln* (5 dpf) | +++<br>(n = 2/2) | +++<br>(n = 1/1) | +++<br>(n = 3/3) | +++<br>(n = 2/2) |
| *olig2* (5 dpf) | +++<br>(n = 3/3) | +++<br>(n = 3/3) | +++<br>(n = 3/3) | +++<br>(n = 3/3) |
| *vglut2a* (5 dpf) | +++<br>(n = 3/3) | +++<br>(n = 1/1) | +++<br>(n = 3/3) | +++<br>(n = 2/2) |

3 or 5-dpf wild-type (WT), *skor1b*, *skor2*, or *skor1b;skor2* mutant larvae were fixed and analyzed by immunostaining with anti-Pvalb7, Ca8, Zebrin II (PC markers) or anti-Vglut1 (GC axonal marker), or by whole mount *in situ* hybridization of riboprobes. Expression levels are indicated by +++++, +++, ++, +, and -. +++++ indicates expressing cells more than those in WT; +++ indicates expression comparable to that in WT; ++ indicates weak expression, + indicates strongly reduced expression; - indicates little or no expression. NA, not appreciable. \*1 Expression was detected in GCs (axons), but not PCs, in the caudal lobe of the cerebellum and rostral hindbrain. \*2 Expression was detected in GCs in the rostral part of the cerebellum (corpus cerebelli).

Although *foxp1b;foxp4* and *skor1b;skor2* mutant larvae showed defects in PC development, expression of *skor1b* and *skor2* was not affected in *foxp1b;foxp4* mutant larvae (Fig. 7Y-AF, Table 2). The expression domains of *foxp1b* and *foxp4* in *skor1b;skor2* double mutant larvae were altered by aberrant differentiation of cerebellar neurons, as described below. In *skor1b;skor2* mutants, *foxp1b* and *foxp4* were observed in the caudal and rostral GC population, respectively (Fig. 8X, AB, Table 3). The expression level of *foxp4* was not strongly affected, while that of *foxp1b* was reduced but not eliminated. These data suggest that *foxp1b/4* and *skor1b/2* are regulated independently of each other in the cerebellum. We further examined *ptf1a* expression in these mutants. *ptf1a* expression was not affected in *skor1b;skor2* mutants (Fig. 8AC-AJ, Table 3), but it was reduced in *foxp1b;foxp4* mutants at 5 dpf but not at 3 dpf (Fig. 8AF, AJ, Table 2). These data suggest that *foxp1b* and *foxp4* function downstream of *ptf1a*, and then function to maintain *ptf1a* expression.

### Skor1b and Skor2 suppress GC fate

We further examined the expression of GC markers in *skor1b;skor2* mutant larvae in more detail. WT, *skor1b* or skor2 single mutant larvae had regions of the cerebellum where Neurod1 expression was absent (Fig. 9A-C, E-G), whereas *skor1b;skor2* mutant larvae did not (Fig. 9D, H). Consistent with this finding, the area of the cerebellum containing Neurod1^+^ GCs was significantly larger in *skor1b*;*skor2* mutant larvae (Fig. 9Q), indicating that immature (Neurod1^+^) GCs increased in the *skor1b*;*skor2* mutant cerebellum. Cell proliferation, indicated by phospho-histone 3, did not increase in the *skor1b*;*skor2* mutant cerebellum (Fig. S5), indicating that increased GCs were not due to an increase in the proliferation of GCs. These data suggest that cells in *skor1b;skor2* mutants that should have differentiated into PCs instead differentiated into Neurod1^+^ immature GCs. We further examined the expression of *cbln12* and *vglut1*, which were reported to be expressed in mature GCs (Bae et al., 2009; Kani et al., 2010; Takeuchi et al., 2017), noting that they did not increase in *skor1b;skor2* mutant larvae (Fig. 9L, P), suggesting that despite an increase in immature GCs, they did not differentiate into mature GCs. To examine the ability of Skor to suppress GC differentiation, RFP (control), or Skor2 together with mCherry, were expressed in GC progenitors in a mosaic manner using *Tg(atoh1c:GAL4FF)* (Kidwell et al., 2018). The expression of Pvalb7 or Neurod1 cells in *atoh1c*^+^-lineage cells expressing transgenes was also examined. When only RFP was expressed, around 60% of cells were Neurod1^+^ cells (Neurod1^-^ cells are likely undifferentiated GCs, Fig. 9U-W, AD). In contrast, when Skor2 and mCherry were co-expressed in *atoh1c*^+^ progenitors, the ratio of the Neurod1^+^ population was significantly reduced (Fig. 9AA-AD). No Pvalb7^+^ cells expressed RFP or Skor2/mCherry (Fig. 9R-T, X-Z). These data indicate that Skor2 can inhibit the differentiation of *atoh1c^+^* GC progenitors to Neurod1^+^ GCs, but Skor2 alone cannot induce the differentiation of *atoh1c^+^*cells to PCs.

## DISCUSSION

### Roles of Ptf1a and Neurog1 in the development of cerebellar neural circuits

Whereas *ptf1a* mutant mice showed a complete loss of GABAergic PCs and INs (Hoshino et al., 2005), *ptf1a* mutant zebrafish showed a partial loss of PCs and INs (Itoh et al., 2020) (Fig. 2), suggesting that the contribution of Ptf1a to the development of PCs and INs differs slightly between mice and zebrafish. Both *ptf1a* and *neurog1* are expressed in the cerebellar VZ in mice and zebrafish (Kani et al., 2010; Lundell et al., 2009) (Fig. 1). Zebrafish *neurog1*;*ptf1a* mutants displayed an almost complete lack of PCs and INs (Fig. 2, Table 1). These data suggest that Ptf1a plays a major role in the development of PCs and INs in zebrafish, whereas Neurog1 functions partially redundantly with Ptf1a in this process. A similar cooperation was observed in the development of crest cells, which were reduced in *ptf1a* mutants and almost absent in *ptf1a;neurog1* mutants (Fig. S2). We previously reported that Ptf1a is essential for the development of IOs in the hindbrain of zebrafish (Itoh et al., 2020). A different dependency of Ptf1a may be explained by overlapping and non-overlapping expression of *ptf1a* and *neurog1* in the rostral (for PC and crest cells) and caudal hindbrain (for IOs) (Fig. 1), as was reported for mice (Yamada et al., 2007). Proneural genes function in neural progenitors to generate various neurons and glial cells in some cases. Lineage tracing revealed that PCs and INs are derived from *ptf1a^+^* neural progenitors (Fig. 4). When considered together, our findings suggest that both PCs and INs in the cerebellum and crest cells in the rostral hindbrain are derived from Ptf1a/Neurog1-expressing neural progenitors in zebrafish.

In addition to PCs and INs, *ptf1a;neurog1* mutants showed reduced expression of *olig2* and *vglut2a* (Fig. 3), which were expressed in ECs in zebrafish cerebellum (Bae et al., 2009; McFarland et al., 2008). Our previous study suggested that *olig2*-expressing ECs were mainly derived from *ptf1a^+^*neural progenitors, but some were derived from *atoh1a^+^* neural progenitors (Kani et al., 2010). Although further lineage tracing of *ptf1a^+^* neural progenitors for ECs is required, the data further support that at least some ECs are derived from Ptf1a/Neurog1-expressing neural progenitors. Furthermore, in *ptf1a;neurog1* mutants, the expression of *atoh1a/b/c* was unaffected at 3 dpf (Fig. S3) but was strongly reduced at 5 dpf (Fig. 3), suggesting that Ptf1a and Neurog1 play a role in maintenance of GC progenitors. It is not clear whether Ptf1a and Neurog1 control the maintenance of GC progenitors cell-autonomously or non-cell autonomously. In mammals, GC progenitors are maintained by Shh produced by PCs (Corrales et al., 2006; Lewis et al., 2004; Wallace, 1999; Wechsler-Reya and Scott, 1999). If a similar mechanism is involved in the maintenance of GC progenitors in zebrafish, the loss of PCs in *ptf1a;neurog1* mutants could lead to their inability to maintain *atoh1a/b/c*-expressing GC progenitors. However, in the zebrafish cerebellum, *shh* is not expressed in PCs and Shh signaling is not activated (Biechl et al., 2016; Chaplin et al., 2010; Hibi et al., 2017). Lineage tracing indicates that at least some GCs were derived from *ptf1a^+^* neural progenitors (Fig. 5). Although a non-cell autonomous role of Ptf1a and Neurog1 in GC development cannot be completely excluded, Ptf1a and Neurog1 likely have a cell-autonomous role in the differentiation of some GCs.

### Does Ptf1a determine GABAergic neural fate?

Loss of function of *ptf1a* and gain of function of *ptf1a* and *atoh1* in mice suggest that Ptf1a and Atoh1 have deterministic roles in the development of GABAergic and glutamatergic neurons, respectively (Hoshino et al., 2005; Pascual et al., 2007; Yamada et al., 2014). However, we found that in zebrafish, *ptf1a*^+^ progenitor cells gave rise to GABAergic PCs, INs, and GCs (Fig. 4, 5, S1). It is possible that *ptf1a* and *atoh1* genes are initially co-expressed in the same neural progenitors in the cerebellum, and these cerebellar neurons are derived from the *ptf1a^+^ atoh1^+^* progenitors. However, PCs, INs, and GCs marked in the lineage tracing experiments were derived from neural progenitors expressing *ptf1a* at 2 dpf (Fig. 4, 5) when the expression regions of *atoh1* genes and *ptf1a* were well separated (Kani et al., 2010). Therefore, at least some of these neurons could be derived from neural progenitors expressing *ptf1a* but not *atoh1* genes. *ptf1a;neurog1* mutants showed an almost complete lack of PCs and INs, but retained GCs at 5 dpf (Fig. 2, 3). Considering that GCs were reported to be mainly derived from *atoh1*^+^ neural progenitors in early-stage larvae (Kani et al., 2010; Kidwell et al., 2018), GCs derived from *ptf1a^+^* neural progenitors are likely to be a minority among GCs. However, our findings indicate that glutamatergic neurons’ GCs and possibly ECs can be generated from *ptf1a^+^* neural progenitors, even if in small numbers, in the zebrafish cerebellum.

The preceding data also suggest that the expression of *ptf1a* in neural progenitors is not sufficient to determine the fate of GABAergic neurons in the zebrafish cerebellum. How can the findings from zebrafish and mice experiments be reconciled? One possibility is that the regulation of downstream genes that determine cell fates by proneural genes is tight in mice, whereas it is more flexible in zebrafish. The expression of GC deterministic genes, such as *neurod1* (Miyata et al., 1999), may be strictly regulated by Atoh1 in mice, but can be regulated by both Atoh1a/b/c and Ptf1a (and Neurog1) in zebrafish. Further analysis is required to understand how proneural genes control the cell fate determination.

### Role of Foxp- and Skor-family transcriptional regulators in PC differentiation

Since *ptf1a^+^* neural progenitors are capable of generating multiple types of cerebellar neurons, there should be factors that determine the cell fate of each type of neuron. We showed that Foxp- and Skor-family transcriptional regulators are expressed in PCs, dependent on Ptf1a and Neurog1 (Fig. 2). The *foxp1b;foxp4* mutant showed a strong reduction of PCs (Fig. 7), and *skor1b;skor2* mutants showed the complete loss of PCs (Fig. 8). Furthermore, Foxp1b, Skor1b, and Skor2 were expressed in differentiating and differentiated PCs (Fig. 6). These data indicate that Foxp1b/4 and Skor1b/2 function downstream of Ptf1a and Neurog1 as key transcriptional regulators during the initial step of PC differentiation. *skor1b* and *skor2* expression was not affected in *foxp1b;foxp4* mutants (Fig. 7). Although the expression region of *foxp1b* and *foxp4* was affected in *skor1b/skor2* mutants due to the aberrant differentiation of cerebellar neurons, their transcripts were retained in the cerebellum (Fig. 8), suggesting that Foxp- and Skor-family proteins function independently to control PC differentiation (Fig. 10).

**Figure 10.**
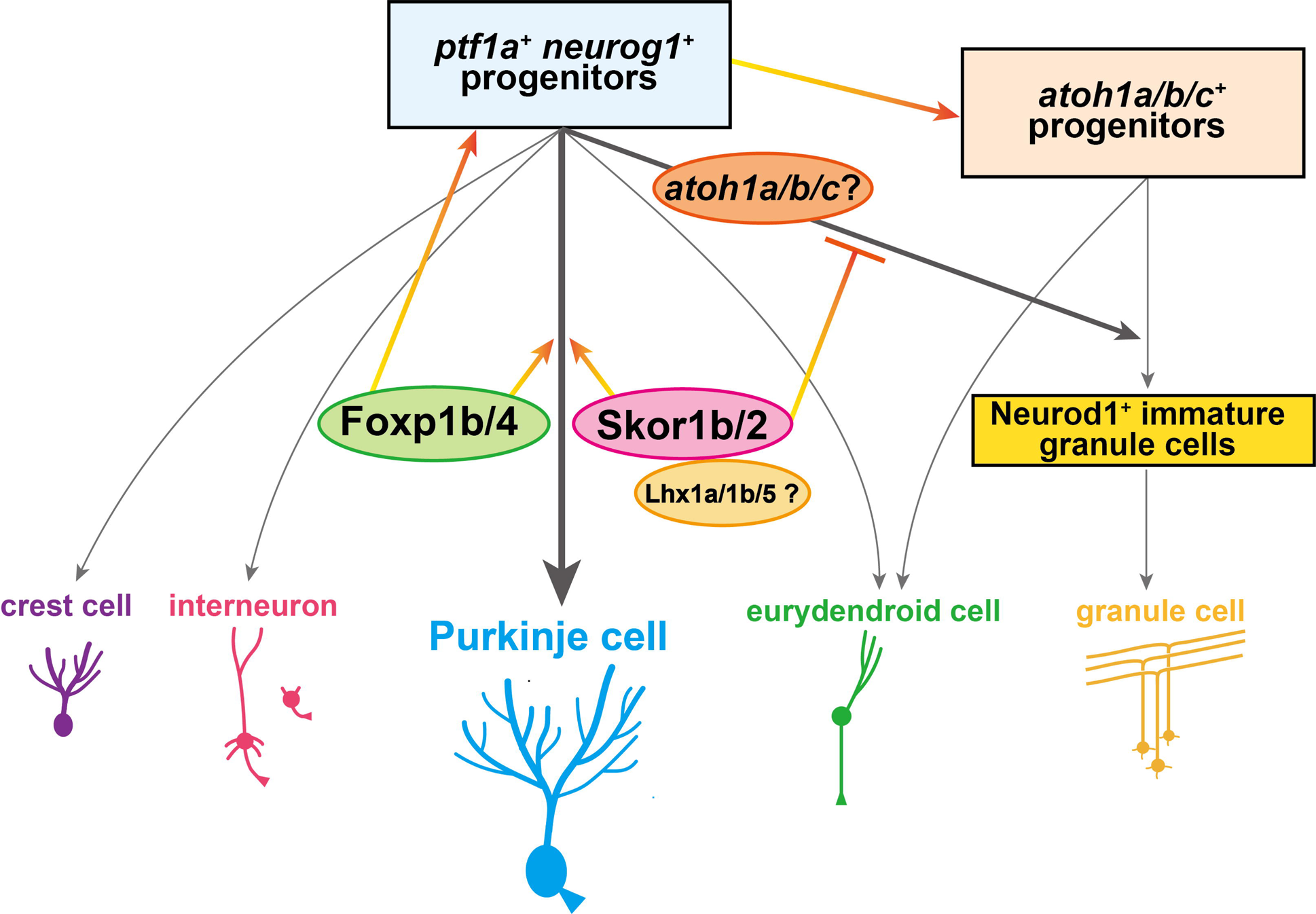
Schematic illustration of a model for neuronal differentiation from Ptf1a/Neurog1-expressing neural progenitors.

Studies of *foxp2*-mutant mice and siRNA-mediated knockdown of *foxp4* in mice revealed that Foxp2 and Foxp4 function in late developmental processes such as cell positioning and dendrite formation (Ferland et al., 2003; Tam et al., 2011; Tanabe et al., 2012). In zebrafish, *foxp1b* and *foxp4* are strongly expressed in PCs while *foxp1a* and *foxp2* are only slightly expressed in PCs (Takeuchi et al., 2017). Thus, zebrafish *foxp1b* may serve the same function as mouse *foxp2*. Although *foxp1b;foxp4* showed a strong reduction of PCs, some PCs remained (Fig. 7). It is possible that the function of *foxp1a* or *foxp2* is partially redundantly with that of *foxp1b* and *foxp4* in PC differentiation. Triple or quadruple zebrafish mutants of *foxp*-family genes should answer this question. *foxp2* is also expressed in IOs in both mice and zebrafish (Fujita and Sugihara, 2012; Itoh et al., 2020). Foxp-family proteins may coordinate differentiation from *ptf1a*^+^ neural progenitors to both PCs and IOs that form the cerebellar neural circuits. In addition to neuronal differentiation, Foxp1b and Foxp4 are involved in the maintenance of *ptf1a^+^* neural progenitors (Fig. 7), indicating that Foxp-family proteins control both the maintenance and differentiation of neural progenitors (Fig. 10). It remains elusive whether Foxp proteins function as transcriptional activators or repressors. Previous studies indicated that Foxp1/2/4 can interact with a component of the NuRD remodeling complex, functioning as transcriptional repressors (Chokas et al., 2010). No increased or ectopic expression of GC genes was observed in the cerebellum of *foxp1b;foxp4* mutants, unlike *skor1b;skor2* mutants (Fig. 7). Further analysis is required to understand the molecular mechanisms of Foxp protein-mediated PC differentiation. Foxp1 is involved in many developmental processes, including specification of motor neuron subtypes in the spinal cord (Dasen et al., 2008; Surmeli et al., 2011). There might be general mechanisms by which Foxp-family proteins control specification from neural progenitors to specific types of neurons.

### Mechanisms of Skor1b- and Skor2-mediated control of PC differentiation

Previous studies on the *skor2* mutant suggested that Skor2 is involved in relatively late development of PCs and the suppression of glutamatergic neuronal genes, but it is dispensable for the initial fate specification of PCs (Nakatani et al., 2014; Wang et al., 2011). We demonstrated that *skor1b;skor2* mutants displayed a complete loss of PCs and instead increase the amount of Neurod1^+^ immature GCs (Fig. 8, 9). Cell proliferation linked to GC proliferation did not increase in *skor1b;skor2* mutants (Fig. S5). *foxp1b* and *foxp4* were normally expressed in differentiating and differentiated PCs but were detected in GCs (Fig. 8). Ectopic expression of *skor2* in GC progenitors reduced the expression of Neurod1 (Fig. 9). These data suggest that in *skor1b;skor2* mutants, cells destined to become PCs differentiated into Neurod1^+^ GCs. Therefore, Skor1b/2 function in the initial step of differentiation from *ptf1a^+^* neural progenitors to suppress differentiation to GCs (Fig. 10). Although Neurod1^+^ GCs increased *skor1b;skor2* mutants, expression of mature GC markers did not increase in these mutants (Fig. 8, 9), indicating that other factors, which possibly function downstream of Atoh1, are required for differentiation of the Neurod1^+^ immature GCs to mature GCs.

It remains elusive whether Skor1b/2 suppress GC fate and thereby secondarily promote PC differentiation, or whether they are also directly involved in PC differentiation independent of GC fate suppression. Mouse Skor2 exhibited transcriptional repression of a reporter in cultured cells (Wang et al., 2011), suggesting that Skor2 directly represses target genes by histone modification. Since the direct binding of Skor family proteins to DNA has not been reported, it is likely that the regulation of gene expression requires transcription factor partners that bind to specific elements of DNA. We screened Skor1b/2 interactors by examining co-immunoprecipitation of Skor1/2 with PC-expressing transcription factors from transfected HEK293T cells and found that zebrafish Skor1b and Skor2 can interact with Lhx-family Lhx1a, Lhx1b, and Lhx5 (there are two genes for Lhx1 in zebrafish, Fig. S6). We generated zebrafish mutants of *lhx1a*, *lhx1b*, and *lhx5* (Fig. S7). As reported in *lhx1;lhx5* mutant mice (Zhao et al., 2007), we found that *lhx1a;lhx5* zebrafish mutants showed a severe reduction of PCs while *lhx1a;lhx1b;lhx5* zebrafish mutants showed a complete loss of PCs (Fig. S8), as did s*kor1b;skor2* mutants. Although Lhx proteins are thought to function as transcriptional activators (Hobert and Westphal, 2000), they may also function with Skor proteins as repressors to repress the expression of GC genes. Alternatively, Skor1b/2 cooperate with Lhx-family proteins to positively promote the expression of some PC genes. The identification of target genes of Skor1b/2 and Lhx1a/1b/5 by chromatin immunoprecipitation (ChIP) should clarify this issue. In any case, Skor- and Lhx-family transcriptional regulators might cooperate to induce PC differentiation and/or suppress GC fate (Fig. 10).

### Gene networks for PC differentiation

In this study, we demonstrate that there are two steps to determine whether cells become PCs or GCs in the cerebellum. In the first step, expression of proneural genes roughly determine cell fate: expression of *atoh1* induces differentiation into GCs while *ptf1a* expression induces the differentiation of PCs. However, expression of proneural genes is not sufficient to determine cell fate. In the second step, Skor-family proteins act as gatekeepers to prevent cells from becoming GCs. Foxp, Skor, and Lhx-family proteins cooperate to promote PC differentiation. The two-step control of PC differentiation ensures that an appropriate number of PCs and GCs are generated to form functional cerebellar neural circuits.

Among *ptf1a^+^* neural progenitors, *foxp1b/4* and *skor1b/2* are only expressed in cells that differentiate into PCs, but not INs, ECs, or GCs. There should be upstream regulators that restrict their expression only to PCs. It was suggested that Olig2 is involved in temporal biasing of PC and IN progenitor fate (Seto et al., 2014). However, *olig2* has different roles in cerebellar neurogenesis, i.e., the development of ECs and oligodendrocytes, in zebrafish (Kani et al., 2010; McFarland et al., 2008). Studies of factors that function upstream and downstream of *foxp*- and *skor*-family genes will provide an understanding of gene networks that control the differentiation of PCs and other cerebellar neurons.

## MATERIALS and METHODS

### Zebrafish strains and genes

The animal work in this study was approved by the Nagoya University Animal Experiment Committee and was conducted in accordance with the Regulations on Animal Experiments at Nagoya University. Wild-type zebrafish with the Oregon AB genetic background were used. For immunohistochemistry and whole-mount *in situ* hybridization, larvae were treated with 0.003% 1-phenyl-2-thiourea (PTU) to inhibit the formation of pigmentation. Zebrafish mutant *ptf1a*^Δ*4*^ (*ptf1a^nub34^*) and *neurog1^hi1059Tg^* were described previously (Golling et al., 2002; Itoh et al., 2020). Transgenic zebrafish *Tg(ptf1a:EGFP)jh1Tg* (Satou et al., 2013), *TgBAC(ptf1a:GAL4-VP16)jh16Tg* (Parsons et al., 2009), *Tg(UAS:RFP)nkuasrfp1aTg*, *TgBAC(neurog1:GFP)nns27Tg* (Satou et al., 2013), *TgBAC(atoh1c:GAL4FF)fh430Tg* (Kidwell et al., 2018), *TgBAC(gad1b:LOXP-DsRed-LOXP-GFP)nns26Tg*, and *TgBAC(slc17ab:LOXP-DsRed-LOXP-GFP)* (Satou et al., 2012) were also described previously. The allele names of the *foxp1b*^Δ*26*^, *foxp4*^Δ*7*^, *skor1b*^Δ*10*^, *skor2*^Δ*8*^, *lhx1a*^Δ*10*^, *lhx1b*^Δ*17*^, and *lhx5*^Δ*10*^ mutants established in this study are designated as *foxp1b^nub89^*, *foxp4^nub90^*, *skor1b^nub91^*, *skor2^nub92^*, *lhx1a^nub93^*, *lhx1b^nub94^*, and *lhx5^nub95^*respectively, in ZFIN (https://zfin.org). The open reading frame (ORF) of *foxp1*, *foxp4*, *skor1b*, and *skor2* mRNAs were isolated by RT-PCR and their sequence information was deposited in DDBJ with the accession numbers LC760469, LC760470, LC760471, and LC760472, respectively. The *skor2* mRNA sequence in a public database (NM_001045421) lacked a region encoding the carboxy-terminal region, so the full ORF of *skor2* was isolated in this study. Zebrafish were maintained at 28°C under a 14-h light and 10-h dark cycle. Embryos and larvae were maintained in embryonic medium (EM) (Westerfield, 2000).

### Establishment of transgenic zebrafish

Establishment of *Tg(5xUAS-hsp70l:mCherry-T2A-CreERT2)* fish will be described elsewhere. This Tg fish harbors transgene(s) containing five repeats of the upstream activation sequence (UAS) and the *hsp70l* promoter (*5xUAS-hsp70l*) (Muto et al., 2017), mCherry, the 2A peptide sequence of *Thosea asigna* virus (TaV), CreERT2 recombinase (Ukita et al., 2009), and the SV40 polyadenylation signal (SV40pAS). To generate *Tg(cbln12:LOXP-TagCFP-LOXP-Kaede)* fish, the TagCFP DNA fragment was amplified from pTagCFP-N (Evrogen) by PCR with the primers 5′-GAAGATCTATAACTTCGTATAGCATACATTATACGAAGTTATACCGGTCGCCACCATGAGCG-3′ and 5′-CCGGAATTCCGGATCCATAACTTCGTATAATGTATGCTATACGAAGTTATACCACAACTAGAATGCAGTG-3′, and subcloned to *BamH*I and *EcoR*I sites of pCS2+ after digestion with *Bgl*II and *EcoR*I (pCS2+lTl). Kaede cDNA from pCS2+Kaede was inserted to *BamH*I and *Xba*I sites of pCS2+lTl-Kaede, which contains SV40pAS. The 2-kpb *cbln12* promoter (Dohaku et al., 2019) and lTl-Kaede-pAS were subcloned to pT2ALR-Dest by NEBuilder (NEB, USA). To generate *Tg(5xUAS-hsp70l:HA-skor2-P2A-mCherry, myl7:mCherry)*, 3xHA (influenza hemagglutinin)-tagged *skor2* cDNAs, the 2A peptide sequence from porcine teschovirus-1 (PTV1), and mCherry cDNA were subcloned into pCS2+, and transferred to the pENTR L5-L2 vector by the BP reaction of the Gateway system. The pENTR L1-R5 plasmid containing *5xUAS-hsp70l* and pENTR L5-L2 containing the *skor2* expression cassette were subcloned into pBleeding Heart (pBH)-R1-R2 (Dohaku et al., 2019), which contains mCherry cDNA and SV40pAS under control of the *myosin*, *light chain 7*, *regulatory* (*myl7*) promoter. To generate *Tg(5xUAS-hsp70l:RFP)*, pENTR L1-L5, which contains *5xUAS-hsp70l*, and pENTR L5-L2, which contains the RFP expression cassette, were subcloned into pT2KALR-Dest (Dohaku et al., 2019). To make transgenic fish, 25 pg of Tol2 plasmid DNA and 25 pg of Tol2 transposase RNA were injected into 1-cell stage WT embryos. The allele names of the Tg line established in this study were designated as *Tg(5xUAS-hsp70l:mCherry-T2A-CreERT2)nub99Tg*, *Tg(cbln12:LOXP-TagCFP-LOXP-Kaede)nub96Tg*, *Tg(5xUAS-hsp70l:HA-skor2-P2A-mCherry*, *myl7:mCherry)nub97Tg*, and *Tg(5xUAS-hsp70l:RFP)nub98Tg* in ZFIN.

### Establishment of zebrafish mutants by the CRISPR/Cas9 system

The gRNA targets were designed by the web software ZiFit Targeter and CRISPRscan (Hwang et al., 2013; Mali et al., 2013; Moreno-Mateos et al., 2015). To generate gRNAs, the following oligonucleotides were used: 5′-TAGGCCGGTGTTCAGAGCACAG-3′ and 5′-AAACCTGTGCTCTGAACACCGG-3′ for *foxp4*^Δ*7*^; 5′-TAGGAGATCCTCAGGCCGCGG-3′ and 5′-AAACCCGCGGCCTGAGGATCT-3′ for *skor1b*^Δ*10*^; 5′-TAGGTTATCATGCCACAGCGC-3′ and 5′-AAACGCGCTGTGGCATGATAA-3′ for *skor2*^Δ*8*^. gRNA and Cas9 mRNA syntheses were performed as previously reported (Nimura et al., 2019). A solution containing 25 ng/μL gRNA and 100 ng/μL Cas9 mRNA or 1000 ng/μL Cas9 protein (ToolGen Inc.) was injected into one-cell-stage embryos using a pneumatic microinjector (PV830, WPI). To establish the *foxp1b* mutant, chemically synthesized crRNAs and tracrRNAs (Fasmac) were used. The following target sequences were selected: 5′-TGGCGTGAGAGGGGCCGTTG-3′. To establish *lhx1a*, *lhx1b*, and *lhx5* mutants, chemically synthesized Alt-R® crRNAs and tracrRNAs, and Cas9 protein (Integrated DNA Technologies, USA) were used. The following target sequences were selected: 5′-GCGAGAGGCCTATATTGGACAGG-3′ for *lhx1a*, 5′-TGAGCGTCTTGGACAGAGCCTGG-3′ for *lhx1b*, and 5′-GTGAGAGGCCCATTCTGGATCGG-3′ for *lhx5*. To prepare the crRNA:tracrRNA Duplex and gRNA, Cas9 RNP Complexes were established as previously reported (Hoshijima et al., 2019). Mutations on the target region were detected by a heteroduplex mobility assay (Ota et al., 2013) and confirmed by sequencing after subcloning the target regions amplified from the mutant genome into pTAC-2 (BioDynamics Laboratory).

### Genotyping

To detect mutations, the following primers were used: 5′-CCCCTCAGTTTACCCCAGA-3′ and 5′-TGAGTAGCGTCTGCGTATGG-3′ (*foxp1b*^Δ*26*^); 5′-TGTTTTAGCCATGTGTCCCACTGA-3′ and 5′-GCTGTTGGTGGTCAGATCGA-3′ (*foxp4*^Δ*7*^); 5′-CCTCTCGGCCTCTCGCTTTGTA-3′ and 5′-CTGGGCATCACCTGTGTGCA-3′ (*skor1b*^Δ*10*^); 5′-AGACATTGTGATGGCAACCCCA-3′ and 5′-CGTAGAGGATGACCTGCCCA-3′ (*skor2*^Δ*8*^); 5′-GGAGCACATCCAAAGACGAT-3′ and 5′-CTTGATGTGCCATGCTCTGT-3′ (*lhx1a*^Δ*10*^); 5′-CAAAACATGGTCCACTGTGC-3′ and 5′-TGCATTTACAGTCACAGCATTG-3′ (*lhx1b*^Δ*17*^); 5′-CGGAATGATGGTGCACTG-3′ and 5′-GTTACACTCGCAGCATTGGA-3′ (*lhx5*^Δ*10*^). To detect WT and *neurog1^hi1059Tg^* mutant alleles, the following three primers were used: 5′-AAAGAAAAGTGGTGGGAAAGCC-3′, 5′-TCGCTTCTCGCTTCTGTTCG-3′, and 5′-GCACAACGTTAGGTATTCACTGTTTG-3′. The WT and *neurog1^hi1059Tg^* mutant alleles gave rise to 412 and 300 bp DNA fragments, respectively.

### Treatment with endoxifen

4 μM endoxifen solution was prepared by adding 0.96 μL of 25 mM endoxifen (Sigma-Aldrich, SML2368) dissolved in DMSO into 6 mL of E3 medium (5 mM NaCl, 0.17 mM KCl, 0.4 mM CaCl_2_, and 0.16 mM MgSO_4_) containing 0.004% PTU. To induce CreERT2-mediated recombination, 2 dpf larvae were treated with the endoxifen solution for 16 h. After washing with E3/PTU medium, larvae were cultivated in this medium until 5 dpf. For the control, DMSO was used instead of 25 mM endoxifen DMSO stock.

### *In situ* hybridization

Whole mount in situ hybridization was performed as previously reported (Bae et al., 2009). Detection of *ptf1a* and *neurog1* was previously described (Bae et al., 2005; Kani et al., 2010). Larvae were hybridized with digoxigenin (DIG)-labeled riboprobes overnight at 65°C and incubated overnight with 1/2000 alkaline phosphatase-conjugated anti-DIG Fab fragment (Roche) at 4°C. BM purple AP substrate (Roche) was used as the alkaline phosphatase substrate. Images were acquired using an Axio-Plan-2 microscope equipped with an AxioCam CCD camera (Zeiss).

### Generation of antibodies and immunohistochemistry

Polyclonal antibodies against Foxp1b, Skor1b, and Skor2 were generated by immunizing rabbits with the synthetic peptides MESIPNQLPAGRDSSC, <u>C</u>IPYANIIRKEKVGTHLNKS, and <u>C</u>HRDYEDDHGTEDML (the underlined C was added to link the peptides covalently with keyhole limpet hemocyanin), respectively. These antibodies were purified using peptide affinity columns that were generated by vinyl polymer resin (TOSOH Bioscience, TOYOPearL AF-Amino-650) and crosslinker m-maleimidobenzoyl-N-hydroxysuccinimide ester (MBS, ThermoFisher Scientific, 22311). For immunostaining, anti-parvalbumin 7 [Pvalb7] (1/1000, mouse monoclonal ascites), anti-carboxy anhydrase 8 [Ca8] (1/100, mouse monoclonal, hybridoma supernatant) (Bae et al., 2009), anti-zebrin II (1/200, mouse monoclonal hybridoma supernatant) (Lannoo et al., 1991), anti-Vglut1 (1/1000, rabbit polyclonal) (Bae et al., 2009), anti-Neurod1 (1/500, mouse monoclonal, hybridoma) (Kani et al., 2010), anti-paired box 2 [Pax2] (1/700, rabbit polyclonal) (BioLegend, 901001), anti-Foxp1b, anti-Skor1b, and anti-Skor2 (1/1000, rabbit polyclonal, affinity purified) were used. CF488A goat anti-mouse IgG (H+L, Biotium, 20018-1), CF568 goat anti-mouse IgG (H+L, Biotium, 20301-1) and CF568 goat anti-rabbit IgG (H+L, Biotium, 20103) were used as the secondary antibodies. Larvae and cryosections were immuno-stained as described previously (Bae et al., 2005; Itoh et al., 2020; Kani et al., 2010). For Skor1b and Skor2 immunostaining, larvae were fixed and treated with acetone 4°C instead of −30°C. An LSM700 confocal laser-scanning microscope was used to obtain fluorescence images. Images were acquired under nearly identical conditions. To show individual cells, confocal optical sections were used (Fig. 6). In Fig. 9 (R-AC), the dynamic range of fluorescence intensity was modified to compensate for differences in the expression of fluorescent proteins and staining conditions.

### Statistics

Data were analyzed using Graphpad PRISM (ver. 5.1 and 6.0) or R software package (ver. 4.2.2).

## Acknowledgements

We thank Shin-ichi Higashijima, Michael J. Parsons, Koichi Kawakami, and the National Bioresource Project for providing the transgenic zebrafish, Masato Kinoshita and Feng Zhang for the hSpCas9 plasmid, Richard R. Behringer and Hiroshi Sasaki for the mCherry-T2A-CreERTe plasmid, Koichi Kawakami for the Tol2-related plasmids, and Kuniyo Kondoh and Yumiko Takayanagi for managing fish mating and care. We also thank the members of the Hibi Laboratory for helpful discussion.

## Competing interests

The authors declare no competing or financial interests.

## Author contributions

Conceptualization: M.H.; Formal analysis: T.I., M.U., S.Y., J.W., A.N.; Data curation: T.S., M.H.; Writing-original draft: T.I., M.H.; Writing-review & editing: T.I., T.S., M.H.; Supervision: M.H.; Funding acquisition: T.S., M.H.

## Funding

This work was supported by the Japan Society for the Promotion of Science KAKENHI (JP15H04376, JP18H02448, and JP22H02631 to M.H., JP18K06333 to T.S.), and Core Research for Evolutional Science and Technology Japan Science and Technology Agency (JPMJCR1753 to M.H.).

## Notes

### Competing Interest Statement

The authors have declared no competing interest.

